# The developing pig respiratory microbiome harbours strains antagonistic to common respiratory pathogens

**DOI:** 10.1101/2023.12.20.572551

**Authors:** Abel A. Vlasblom, Birgitta Duim, Shriram Patel, Roosmarijn E. C. Luiken, Daniel Crespo-Piazuelo, Julia Eckenberger, Chloe E. Huseyin, Peadar G. Lawlor, Christian Elend, Jaap A. Wagenaar, Marcus J. Claesson, Aldert L. Zomer

**Affiliations:** Division of Infectious Diseases and Immunology, Faculty of Veterinary Medicine, Utrecht University, the Netherlands; WHO Collaborating Centre for Reference and Research on Campylobacter and Antimicrobial Resistance from a One Health Perspective / WOAH Reference Laboratory for Campylobacteriosis, Utrecht, the Netherlands; School of Microbiology and APC Microbiome Ireland, University College Cork, Cork, T12YT20, Ireland; SeqBiome Ltd., County Cork, Ireland; Teagasc, Pig Development Department, Animal & Grassland Research & Innovation Centre, Moorepark, Fermoy, Co. Cork, P61 C996, Ireland; EW Nutrition Innovation GmbH & Co.KG, 50829 Cologne, Germany; Wageningen Bioveterinary Research, Lelystad, the Netherlands

## Abstract

In the global efforts to combat antimicrobial resistance and reduce antimicrobial use in pig production, there is a continuous search for methods to prevent and/or treat infections. Within this scope, we explored the relationship between the developing piglet nasal microbiome and (zoonotic) bacterial pathogens from birth until ten weeks of life. The nasal microbiome of 54 pigs was longitudinally studied over 16 time-points on nine farms in three European countries (Germany, Ireland, and the Netherlands) using amplicon sequencing targeting the V3-V4 16S rRNA region as well as the *tuf* gene for its *Staphylococcal* discrimination power. The piglets’ age, the farm, and the litter affected the nasal microbiome, with piglets’ age explaining 19% of the variation in microbial composition between samples. Stabilization of the microbiome occurred around two weeks post-weaning. Notably, while opportunistic pathogens were ubiquitously present, they did not cause disease. The piglet nasal microbiome often carried species associated with gut, skin, or vagina, which suggests that contact with the vaginal and faecal microbiomes shape the piglet nasal microbiome. We identified bacterial Co-Abundance Groups (CAGs) of species that were present in the nasal microbiomes in all three countries over time. Anticorrelation between these species and known bacterial pathogens identified strains that might be exploited for pathogen reduction. Further experimental evidence is required to confirm these findings. Overall, this study advances our understanding of the longitudinal development and factors influencing the piglet nasal microbiome, providing insights into its role in health and disease.

**Importance:** Our study on longitudinal analysis of the developing nasal microbiota of piglets in farms in three European countries showed consistent microbiome compositions and that colonization of porcine pathogens occurred in relation with anticorrelating species. These findings enhance our knowledge of co-colonizing species in the nasal cavity, and the identified microbial interactions can be explored for the development of interventions to control pathogens in porcine husbandry.

## Introduction

In pig farming, bacterial respiratory and systemic infections can be detrimental to health and welfare, and increase cost and antimicrobial use (1, 2). There is a continuous search for (biological) interventions, such as probiotics and competitive exclusion strategies, to prevent and treat infections (3–5).

The impact of the respiratory microbiome on piglet respiratory or systemic infections is an emerging field (6–8). Differences in (upper) airway microbiome composition between healthy and diseased pigs (9–15), between livestock associated methicillin-resistant *Staphylococcus aureus* (LA-MRSA) carriers and non-carriers (7, 16, 17) and due to the farm environment such as gaseous ammonia concentrations (18) have been described, often in cross-sectional studies. These studies have identified species associated with disease, but longitudinal development of the pig nasal microbiome (PNM) in relation with bacterial pathogens is poorly understood. By elucidating the PNM’s intricacies, we will obtain insights that might improve animal health through microbiome modulation.

Opportunistic bacterial pathogens, including those part of the Porcine Respiratory Disease Complex (PRDC) (1, 19), such as *Actinobacillus pleuropneumoniae* (20), *Actinobacillus suis* (21), *Trueperella pyogenes* (22), *Bordetella bronchiseptica* (20), *Glaesserella parasuis* (11), *Klebsiella pneumoniae* (23), *Mannheimia varigena* (24), *Mycoplasma hyopneumoniae* (20), *Mycoplasma hyorhinis* (25), *Pasteurella multocida* (26), and *Streptococcus suis* (27) are often present in the porcine upper respiratory tract. In addition, potential zoonotic pathogens like LA-MRSA reside in the respiratory tract, and zoonotic spill-over of LA-MRSA occurs through occupational exposure to livestock, likely through animal contact or dust (28–30).

To elucidate trends in the development of the PNM in relation to pathogens, 54 piglets were sampled (nasal swabs) across nine farms in three European countries from birth until 70 days of age for *tuf* and 16S rRNA gene amplicon sequencing. Correlation network analysis identified co-abundance groups (CAGs), and species which displayed anti- and co-correlation to bacterial pathogens.

## Results

### Cohort characteristic and sampling summary

Nasal swabs were obtained from 54 piglets born to 27 sows across nine farms equally distributed in three countries (Table 1). We sampled from birth up to 10 weeks of age; daily during the first week of life, as our previous work showed rapid microbiome developed during this period (7), and weekly thereafter. The 16S rRNA was sequenced from all samples (n = 813; Table 1). Furthermore, samples from two of the three litters per farm were additionally *tuf* amplicon sequenced (n = 538) for improved *Staphylococcus* resolution. Due to sequencing failure 21 samples were lost (dark grey values in Table 1). Another 30 samples were excluded due to doxycycline treatment after timepoint 27 in farm NLD3 (light grey 0 values in Table 1).

**Table 1.**
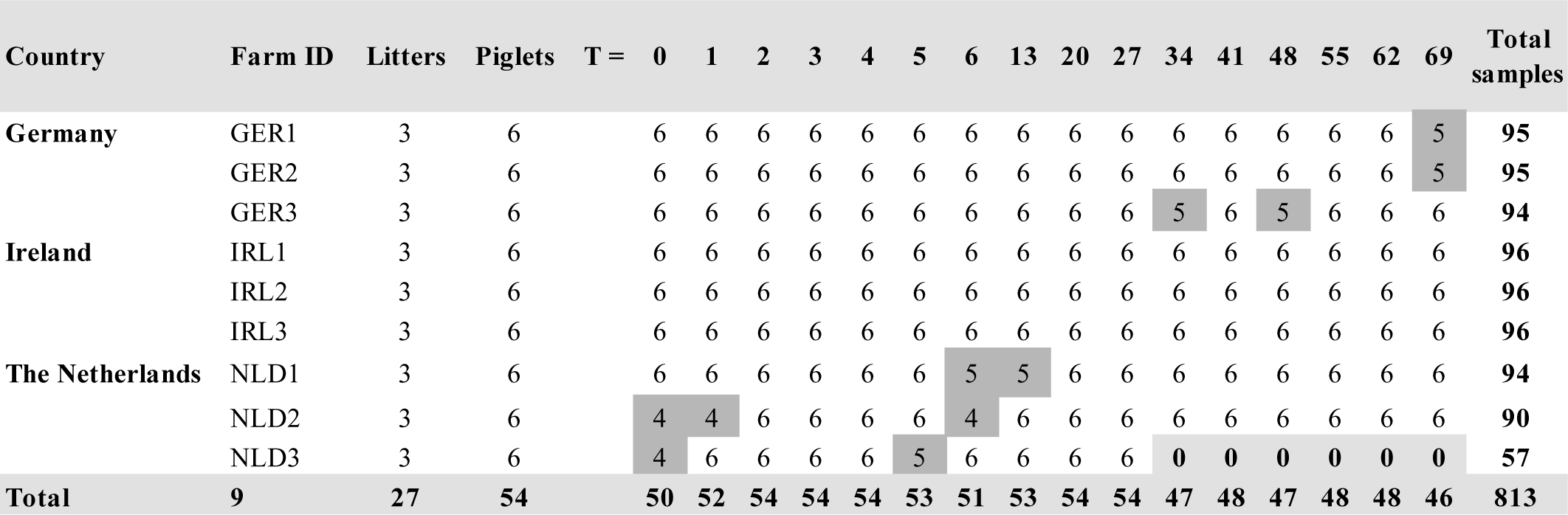
Distribution of nasal swab samples used for amplicon sequencing. Timepoints are depicted in days (0–69) and represent the piglet’s age. The numbers under the timepoints represent the nasal swabs samples collected.

### Summary of the sequencing results

Sequencing generated 32 and 15 million raw 16S rRNA and *tuf* gene reads, respectively. Microbiome analysis was performed on 17 and 11 million curated reads with a mean count of 19,812 ± 5,298 standard deviation (SD) and 18,226 ± 5,782 SD reads/sample for 16S rRNA and *tuf*, respectively (Supplementary Figures S.1A, S.1B and S2.A, S2.B). For 16S rRNA, 11 negative-controls had a mean read count of 1204 with 3 of the 11 negative-controls containing ∼4000 reads. For *tuf*, seven negative controls were sequenced with each of them have very few reads in them (<20 reads). Overall, 10,088 unique amplicon sequence variants (ASVs) were identified in the 16S rRNA dataset, 1430 of which were detected as potential contaminant. For *tuf*, 4,038 unique ASVs were generated with no potential contaminants. In the 16S rRNA dataset, the phylum *Proteobacteria*, with a 65.4% mean relative abundance (MRA), was most prevalent followed by *Firmicutes* (20.8% MRA), and *Bacteroidetes* (7.2% MRA) (Figure S7.A). At genus level, *Moraxella* (44.9%) had the highest MRA followed by *Streptococcus*, *Mannheimia, Rothia*, and *Actinobacillus* all at ∼5% MRA. For *tuf*, *Proteobacteria* and *Firmicutes* made up around 100% (59.5% and 39.7%, respectively) of the detected phyla (Figure S7.B), and the largest genera were *Moraxella* (59.0%) and *Streptococcus* (16.8%).

### Bacterial diversity changes in the porcine nasal microbiome over time

The species diversity, estimated with the Shannon index was higher at birth (Figure 1) for the *tuf* and 16S rRNA datasets with a lower overall richness for *tuf*. We observed a diversity decrease in the first week of life (T=0-6) which was most pronounced in the Netherlands, followed by a diversity increase at T=13 in the Dutch farms. This increase subsided before week five (T=34). There was no significant effect of weaning (around T=27) in the 16S rRNA dataset. For *tuf* there was a significant effect (Welch two sample t-test, p 0.035 and p 0.011: α 0.05) between the average Shannon diversity of week two and three (T=13 and T=20) versus week four (T=27), with week four samples having lower average Shannon diversity than samples before week three.

**Figure 1.**
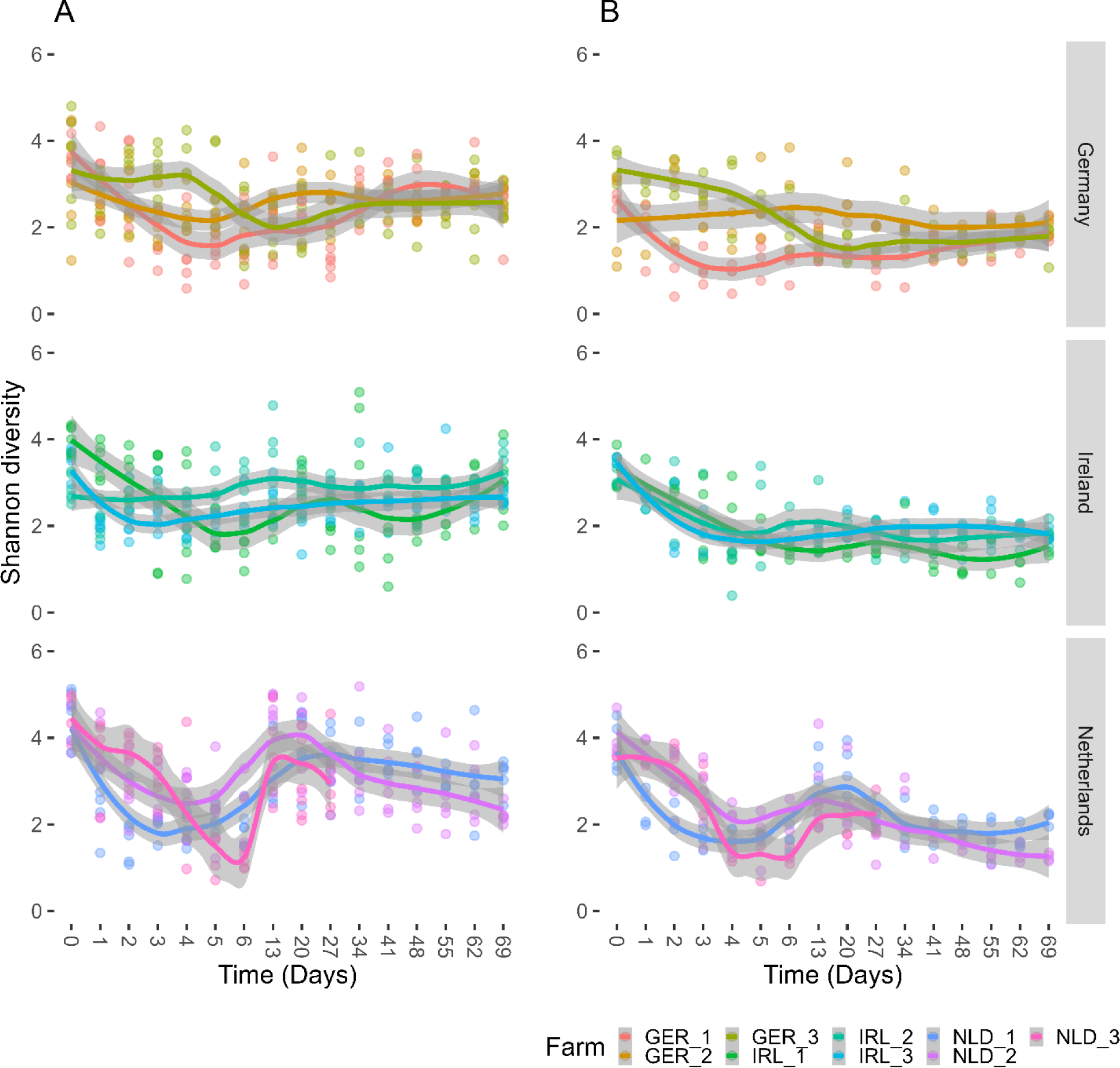
Longitudinal changes in the piglet nasal community Shannon diversity measured for **A:** 16S rRNA and **B**: *tuf* gene amplicon sequencing. X-axis: relative sample time (days), timepoint 0 represents the piglets’ day of birth. Y-axis: Shannon diversity index. Farms are presented by country, and colour coded for identification. The dots represent the samples, lines display the predicted mean and the grey boundaries the standard error.

### Beta diversity displays country independent microbiome development

To investigate compositional changes in the piglet nasal microbiota over time, beta diversity was assessed via principal component analysis (PCA) based on Aitchison distances (Figure 2). Samples from day 0, 1, 2 are visibly distinctly clustered from samples taken later than day 34 (one-week post weaning). These observations were consistent across all three countries and validated by permutational analysis of variance (PERMANOVA), which showed that time (age of the piglet) explained most of the variation in both the 16S rRNA and *tuf* datasets (19.24% and 24.73%, respectively; Table 2). The 16S rRNA Shannon diversity of the piglet nasal swabs trends with the PC2-axis of the ordination plots (higher diversity at the bottom of the graph). For *tuf* the diversity lowers along the PC1-axis, being highest directly after birth.

**Figure 2.**
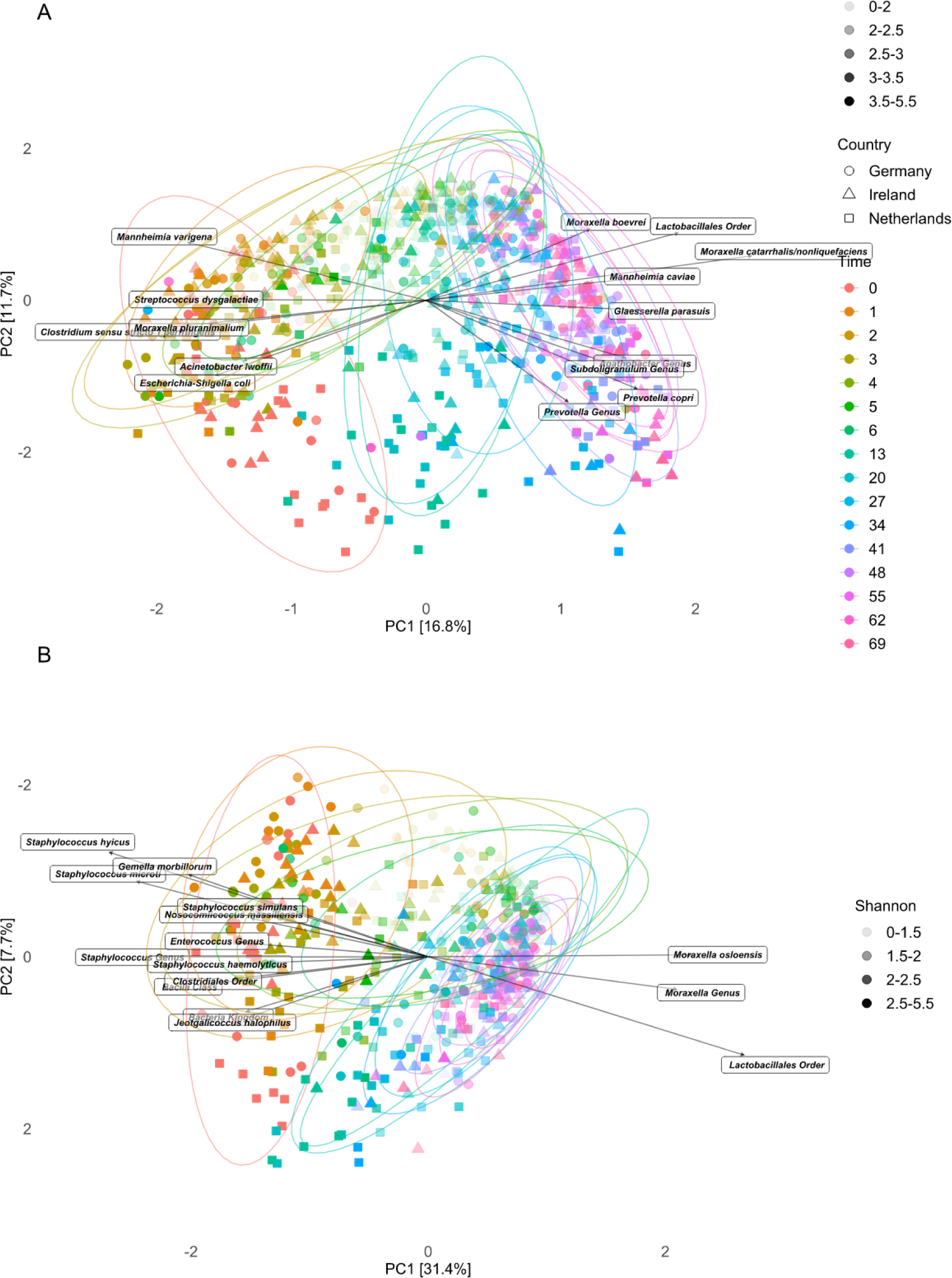
Principal Component Analyses show compositional differences and similarities (beta diversity) in the nasal microbiomes of piglets over time in the 16S rRNA data (**A**) and the *tuf* data (**B**). Samples were coloured by age of the piglets in days (0–69) and given shape by country (Germany, Ireland, and the Netherlands). The arrows show the top 15 ordination driving taxa. Transparency of the plotted points indicate the Shannon diversity of each sample. The 16S rRNA and the *tuf* figures have different transparencies due to the lower Shannon diversity for *tuf*.

**Table 2.** PERMANOVA outputs for the 16S rRNA and *tuf* datasets with time, sex, and the nested factor country, farm, sow, piglet as factors explaining variation.

| 16S rRNA |  |  |  |  | *tuf* |  |  |  |
| --- | --- | --- | --- | --- | --- | --- | --- | --- |
| Factor | Df | SumOfSqs | R<sup>2</sup> | Pr (>F) | Df | SumOfSqs | R<sup>2</sup> | Pr (>F) |
| Time | 15 | 132473.46 | 0.1924 | 0.0002* | 15 | 29408.61 | 0.2473 | 0.0002* |
| Sex | 1 | 1794.97 | 0.0026 | 0.0008* | 1 | 462 | 0.0039 | 0.0030* |
| Country | 2 | 19065.64 | 0.0277 | 0.0002* | 2 | 3972.26 | 0.0334 | 0.0002* |
| Country:<br>farm | 6 | 36632.69 | 0.0532 | 0.0002* | 6 | 8678.21 | 0.0730 | 0.0002* |
| Country:<br>farm: sow:<br>piglet | 18 | 25908.57 | 0.0376 | 0.0002* | 9 | 4159.9 | 0.0350 | 0.0002* |
| Country:<br>farm: sow:<br>piglet | 26 | 14908.61 | 0.0216 | 0.8660 | 17 | 2122.51 | 0.0178 | 0.8676 |
| Residual | 744 | 457918.93 | 0.6649 | NA | 487 | 70125.58 | 0.5896 | NA |
| Total | 812 | 688702.86 | 1 | NA | 537 | 118929.1 | 1 | NA |

\* Represents a significant adjusted P-value ( $\alpha < 0.05$ )

The arrows in the 16S rRNA PCA plot indicate that shifts along PC2 towards the top were driven by species from the genera *Moraxella* and *Mannheimia*, while *Prevotella* and *Escherichia* were drivers of the ordination shift to the bottom samples. Shifts along PC1 were driven by the genera *Clostridium*, *Moraxella*, *Streptococcus* (early samples), *Moraxella*, *Glaesserella*, and *Mannheimia* (late samples) (Figure 2). For *tuf,* 12 of the top 15 ordination driving taxa were attributed to samples from earlier timepoints. *Moraxella osloensis*, *Moraxella* genus, and the *Lactobacillales* order drove the shift along PC1 towards later timepoint clusters.

We sought to identify factors that explained most of the variation in the beta diversity, apart from the factor time (19.24% and 24.73%). We therefore tested the explained variance of the hierarchical levels and found that all levels explained some of the variance, but farm explained a small majority (5.3% and 7.3%, Table 2). Sex of piglets had a significant but small (nested PERMANOVA; 0.26% and 0.39%; p = 0.008 and 0.003) effect on the variation in both datasets.

### Temporal trends in relative abundances at species level taxonomy

Individual farm-level differences were observed among the top 25 most abundant taxa at species level (Figure S.5). However, PNM trends between the countries seemed to be conserved (Figure 3). Sample composition changed with time, where earlier samples had a different composition to samples later in life. Generally, the microbiome was stable after day 34. For 16S rRNA, the most abundant species were members of the genus *Moraxella* which often accounted for half of the species abundance. During the first week after birth, *M. boevrei,* together with *M. bovoculi* and *M. porci,* were the most abundant. Towards the end of the first week the species *M. catarrhalis/nonliquefaciens*, and an unknown species of the genus *Moraxella*, became more prevalent. The species *M. porci* was consistently observed throughout all timepoints (prevalence bar, Figure 3C), with its highest abundance beginning at week one and after day 48 with a pronounced larger proportion in the later German samples (Figure 3A). *Streptococcus suis* was prevalent throughout with a low mean abundance. During the first week, we prevalently observed the species *Rothia nasimurium* and *Mannheimia varigena*, and species from the genera *Clostridium* and *Escherichia*, both gastrointestinal tract associated genera, were abundant (31). From the end of week 1 to week 10 we observed an increasing fraction of *Actinobacillus indolicus* (most pronounced in Ireland)*, Bergeyella zoohelcum* and *Glaesserella parasuis*. These species together with the various *Moraxella* species, comprised on average half of the averaged relative abundances in week 1-10. Weaning, which occurred around day 27, had a small effect on the most abundant taxa in all three countries. Compositional differences between week one (T=7) and week two (T=13) were observed, especially in the Dutch samples. In week two (T=13), fewer of the top 25 most abundant species were present. This is most clearly observed in Dutch farm NLD2 (Figure S.5A).

**Figure 3.**
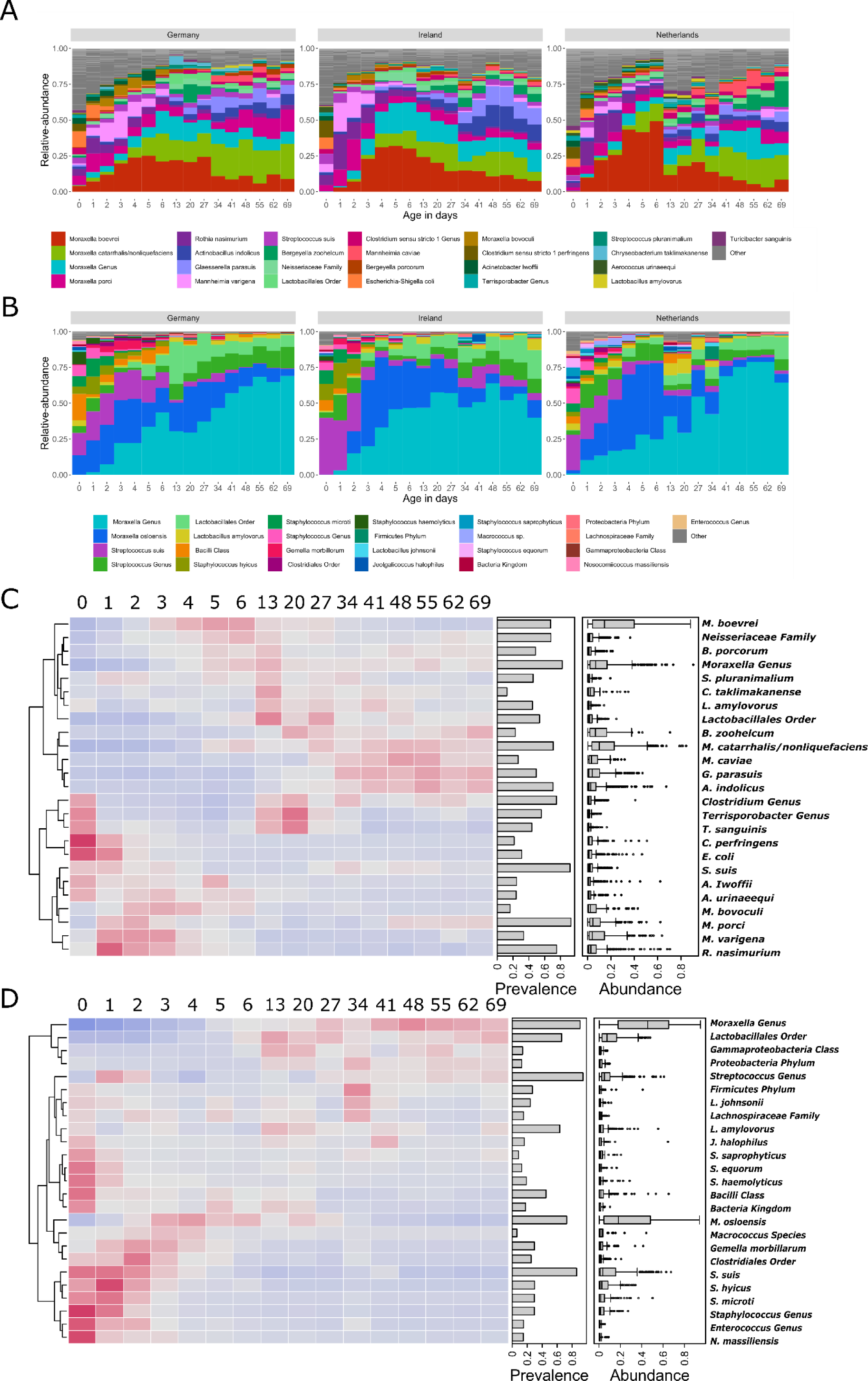
The top 25 most abundant taxa at species level in the 16S rRNA (**A**) and *tuf* (**B**) data in all samples per country. The piglet samples are grouped by age of the piglet in days. The heatmaps show, for each of the 25 top species, at which timepoint each species had its highest relative abundance in the 16S rRNA (**C**) and *tuf* (**D**) data with red indicating a higher relative abundance and blue indicating lower abundance of that taxon. The bars to the right of the heatmap show the proportion of species prevalence over all samples, and boxplots show the distribution of its relative abundance over all samples.

A lower number of taxa for *tuf*, an amplicon known for a high *Firmicutes* resolution especially for *Staphylococcus*, *Streptococcus*, and *Enterococcus* species (32–34), were observed. In the *tuf* dataset, the *Moraxella* genus was by far the most abundant taxon, specifically an unclassified member of the genus *Moraxella* (Figure 3B). The second abundant *Moraxella* species was *M. osloensis,* which could not be discriminated by our 16S rRNA gene sequencing and was abundant until weaning (T=27). The species *S. suis* had its highest relative abundance in the first days of life but was detected throughout the sampling period (high prevalence bar, Figure 3D). Around week two, the *Streptococcus* genus and *Lactobacilliales* order, both without better taxonomic resolution, became more abundant.

Like 16S rRNA, the composition of the Dutch *tuf* samples changed between week one (T=6) and week two (T=13). The top 25 taxa of all samples at species level comprised 90-100% of the relative abundance. After weaning, 4 to 5 species made up 90% of the relative abundance, illustrating the low number of taxa represented by *tuf*.

### Temporal trends of bacterial pathogens in the piglet nasal microbiota

The relative abundances in PNM showed that of the 12 pig pathogens studied, specified in Table 3, nine were detected using 16S rRNA, and two using *tuf* (*S. aureus* and *S. suis*).

**Table 3.** Presence of bacterial pathogens in the 16S rRNA dataset.

| Species | Prevalence<br>in samples<br>(N / 813) | Average relative<br>abundance when<br>detected | Average relative<br>abundance<br>overall | Maximum relative<br>abundance in a<br>sample |
| --- | --- | --- | --- | --- |
| *Actinobacillus pleuropneumoniae* | 136 | 1.96% | 0.33% | 56.43% |
| *Actinobacillus suis* | ND* | - | - | - |
| *Bordetella bronchiseptica* | ND* | - | - | - |
| *Glaesserella parasuis* | 500 | 6.20% | 3.81% | 47.00% |
| *Klebsiella pneumoniae* | 69 | 0.87% | 0.07% | 6.69% |
| *Mannheimia varigena* | 365 | 7.50% | 3.40% | 63.71% |
| *Mycoplasma hyopneumoniae* | ND* | - | - | - |
| *Mycoplasma hyorhinis* | 196 | 2.12% | 0.51% | 51.45% |
| *Pasteurella multocida* | 84 | 1.02% | 0.11% | 17.12% |
| *Staphylococcus aureus* | 11 | 0.29% | < 0.01% | 0.69% |
| *Streptococcus suis* | 804 | 3.15% | 3.11% | 25.43% |
| *Trueperella pyogenes* | 55 | 0.29% | 0.02% | 3.20% |

\* These species have not been detected in the sequencing data.

In the 813 samples, *S. suis* was most prevalent (804/813) followed by *G. parasuis* (500/813), and *M. varigena* (365/813). These species also had the highest MRA of the 12 pathogenic species, around 3% across samples. *M. varigena* had the highest MRA of 7.5%. The maximum relative abundance of *M. varigena, A. pleuropneumoniae*, *G. parasuis*, and *M. hyorhinis* was around 50%, and *S. aureus* had the lowest maximum abundance at around 0.7. The species *Bordetella bronchiseptica* was not identified, but ASVs assigned to the *Bordetella* genus were present. *T. pyogenes, S. aureus,* and *K. pneumoniae* were rarely detected, with a peak of *T. pyogenes* in farm NLD1 samples containing a relative abundance > 0.1*%* for a period from week 3 till week 8 (Figure 4), and *K. pneumoniae* was detected in the three German farms and in farm IRL3.

**Figure 4.**
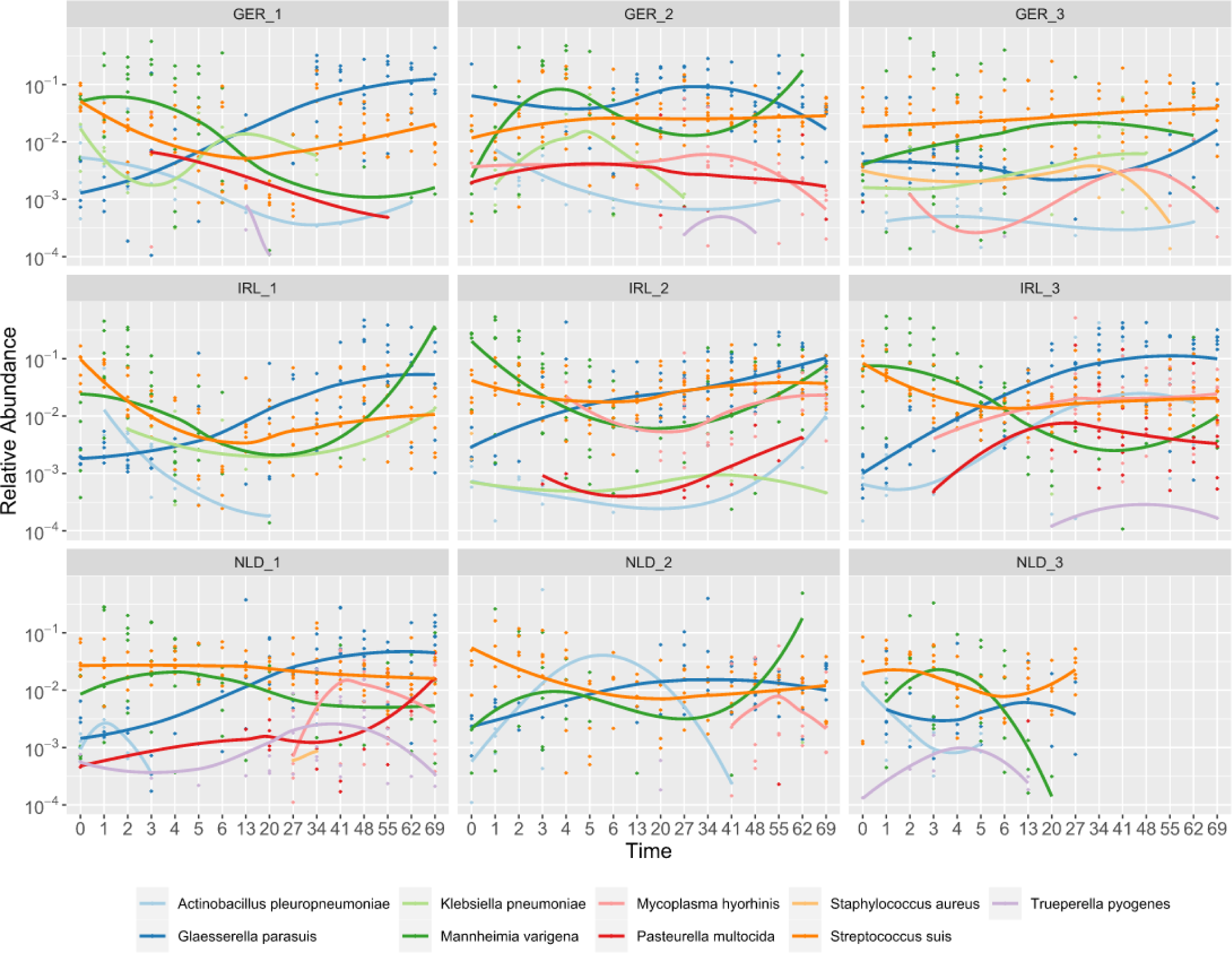
Log-transformed relative abundances (> 0.0001) for 16S rRNA over time of the nine detected pig pathogens. Facetted per farm. Lines indicate trends in abundance within a farm.

To investigate whether there were temporal trends in the presence of pathogens, we plotted the relative abundance of the pathogens investigated for 16S rRNA (Figure 4) and *tuf* (Figure S7) by farm. As expected from the high prevalence of *S. suis* (Table 2), this taxon was consistently detected across the studied farms. *A. pleuropneumoniae* was mostly detected in the first week except for farm IRL3 where it increased in abundance after week two. *G. parasuis* was detected throughout the sampling period, generally with higher abundance after week two (except for farm GER3 where it was only detected in the first week).

We observed different profiles of pathogen presence between farms. For example, in farms GER1 and IRL1 hardly any *M. hyorhinis* or *P. multocida* were detected. In contrast, co-occurrence of these species was observed on farm IRL2, IRL3 and NLD1. *M. hyorhinis* was prevalent in five of the nine farms and mostly after week two.

### Co- and anti-correlation of bacterial species in the piglet nasal microbiome

Positive and negative associations between the top ∼100 species were determined per country after removing species with low prevalence. Species, having a SparCC correlation index of smaller than - 0.2 and bigger than 0.2 per country, were included for clustering using a Markov cluster algorithm (MCL). We observed seven distinct co-abundance groups (CAGs) for 16S rRNA and two *tuf*- CAGs. 16S-CAG1 was the largest with 15 members, followed by 16S-CAG2 with 13 members. We further established two *tuf*-CAGs of similar size (7 and 8 taxa) (Supplementary Table 1).

The SparCC inferred interactions of the species were visualised in a network (Figure 5). Between the members of 16S-CAG1 and 16S-CAG2, exclusively negative interactions were observed. Members of 16S-CAG3 and 16S-CAG4 shared positive interactions between them and negative interactions with members of the other 16S-CAGs. The smaller 16S-CAGs (16S-CAG5, 16S-CAG6, and 16S-CAG7) had positive associations with members of 16S-CAG2.

**Figure 5.**
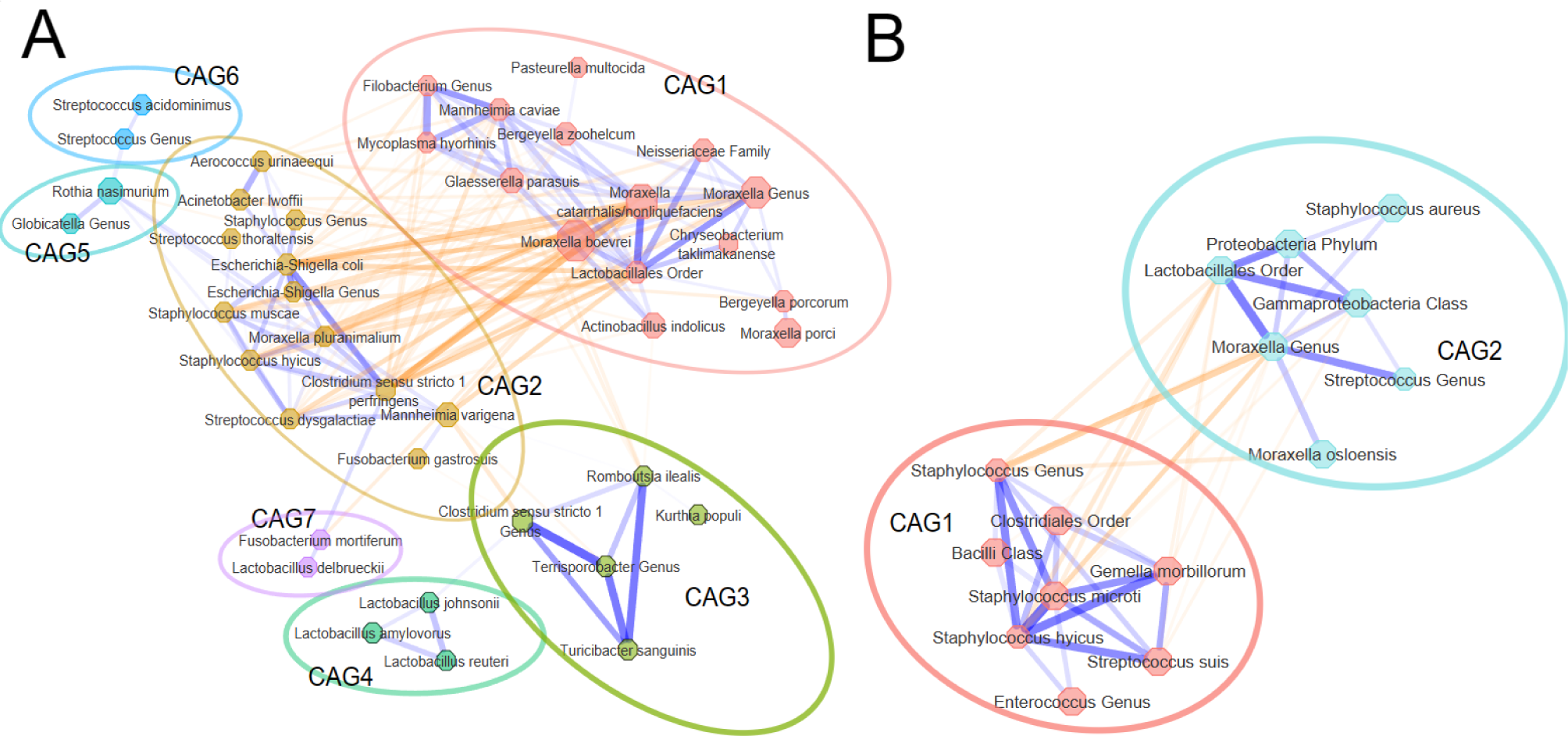
Network of co- and anti-correlating species for 16S rRNA (**A**) and *tuf* (**B**). Line colour orange indicates negative association and blue indicates positive association. A thicker line indicates a stronger interaction. The larger the node size indicates greater abundance in the dataset. Species are coloured according to their CAGs.

### Temporal succession of taxa could underlie the co-abundance grouping

Time explained most variability in microbial composition. To observe time effects on CAGs, the average of the summed relative abundances per CAG (Figure 6) were plotted over time. In all three countries similar trends were observed; 16S-CAG1 increased in abundance from day one and remained dominant up to day 70 of life. Inversely, the abundance of 16S-CAG2 was highest at birth and decreased from the first day, with some fluctuations in abundance later in life. Similarly, *tuf*-CAG1 was succeeded by *tuf*-CAG2 after day one, a trend that was again evident in all three countries.

**Figure 6.**
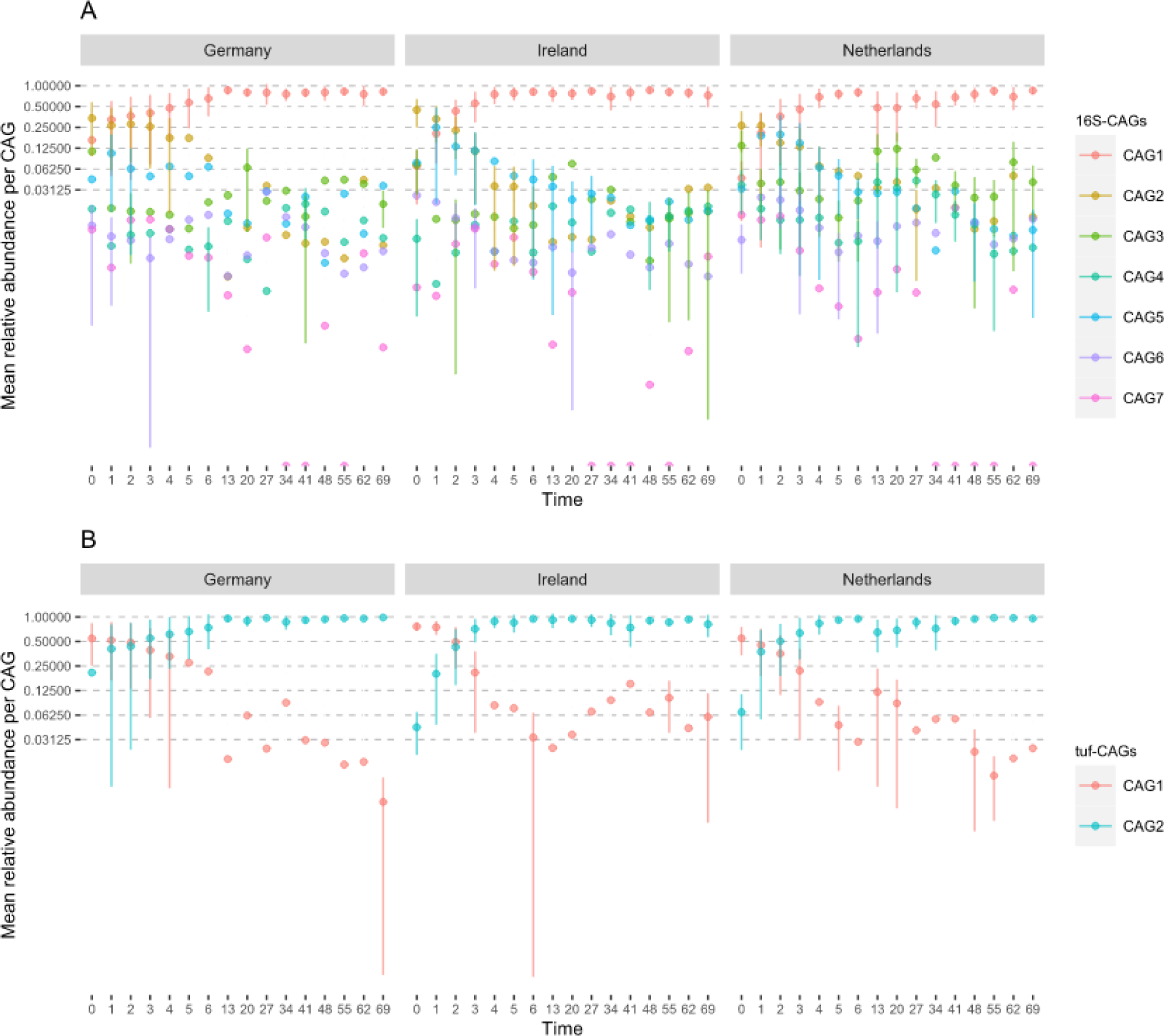
Average relative abundances of the seven 16S-CAGs (**A**) and two *tuf*-CAGs (**B**) by time, per country, error bars represent the standard deviation in relative abundance between samples.

16S-CAG3 had a minor peak around week two of life (two weeks prior to weaning). This effect was more evident in the unaveraged plot of 16S-CAG3 (Supplementary Figure 3). The 16S-CAGs other than the 16S-CAGs 1&2 had a generally low relative abundance.

### Candidate probiotic species that are anti-correlated with respiratory pathogens

We utilised the SparCC analysis of the 16S rRNA and *tuf* datasets to infer taxa negatively associated with airway pathogens over all three countries, displayed in Table 4. Porcine pathogens of 16S-CAG1 (*G. parasuis*, *M. hyorhinis*, and *P. multocida*) showed inverse correlations with members of 16S-CAG2, 16S-CAG3 and 16S-CAG5. *M. varigena*, a pathogen of 16S-CAG2, was negatively correlated with members of 16S-CAG1, 16S-CAG3, and 16S-CAG4 (Table 4). These anticorrelating taxa could be explored for probiotic potential.

**Table 4.** Significant anticorrelating taxa with common airway pathogens. The anticorrelation is based on the average SparCC r value over Germany, Ireland, and the Netherlands. All taxa displayed negative correlations (p < 0.0001) with an average r < −0.2 over the three countries.

| Pathogen | Taxon anticorrelated | Correlation | Dataset | CAG | Pathogen | Taxon anticorrelated | Correlation | Dataset | CAG |
| --- | --- | --- | --- | --- | --- | --- | --- | --- | --- |
| *Glaesserella parasuis* **CAG1** |  |  |  |  | *Mycoplasma hyorhinis* **CAG1** |  |  |  |  |
|  | *Clostridium sensu stricto 1 perfringens* | -0.35 | 16S | CAG2 |  | *Clostridium sensu stricto 1 perfringens* | -0.33 | 16S | CAG2 |
|  | *Streptococcus dysgalactiae* | -0.3 | 16S | CAG2 |  | *Escherichia-Shigella coli* | -0.30 | 16S | CAG2 |
|  | *Romboutsia ilealis* | -0.3 | 16S | CAG3 |  | *Romboutsia ilealis* | -0.28 | 16S | CAG3 |
|  | *Turicibacter sanguinis* | -0.28 | 16S | CAG3 |  | *Aerococcus urinaeequi* | -0.28 | 16S | CAG2 |
|  | *Staphylococcus hyicus* | -0.27 | 16S | CAG2 |  | *Acinetobacter lwoffii* | -0.26 | 16S | CAG2 |
|  | *Mannheimia varigena* | -0.26 | 16S | CAG2 |  | *Staphylococcus hyicus* | -0.25 | 16S | CAG2 |
|  | *Acinetobacter lwoffii* | -0.26 | 16S | CAG2 |  | *Streptococcus dysgalactiae* | -0.25 | 16S | CAG2 |
|  | *Rothia nasimurium* | -0.26 | 16S | CAG5 |  | *Moraxella pluranimalium* | -0.25 | 16S | CAG2 |
|  | *Aerococcus urinaeequi* | -0.26 | 16S | CAG2 |  | *Turicibacter sanguinis* | -0.23 | 16S | CAG3 |
|  | *Moraxella pluranimalium* | -0.24 | 16S | CAG2 |  | *Staphylococcus muscae* | -0.20 | 16S | CAG2 |
|  | *Escherichia-Shigella coli* | -0.23 | 16S | CAG2 |  |  |  |  |  |
|  | *Acinetobacter* Genus | -0.23 | 16S | - | *Pasteurella multocida* **CAG1** |  |  |  |  |
|  | *Staphylococcus muscae* | -0.22 | 16S | CAG2 |  | *Moraxella pluranimalium* | -0.21 | 16S | CAG2 |
|  |  |  |  |  |  | *Clostridium sensu stricto 1 perfringens* | -0.20 | 16S | CAG2 |
| *Mannheimia varigena* **CAG2** |  |  |  |  |  |  |  |  |  |
|  | *Clostridium sensu stricto 1* Genus | -0.4 | 16S | CAG3 | *Staphylococcus aureus* *tuf* **CAG2** |  |  |  |  |
|  | *Moraxella catarrhalis/nonliquefaciens* | -0.34 | 16S | CAG1 |  | *Bacilli* Class | -0.23 | *tuf* |  |
|  | *Lactobacillales* Order | -0.31 | 16S | CAG1 |  |  |  |  |  |
|  | *Terrisporobacter* Genus | -0.27 | 16S | CAG3 | *Streptococcus suis* *tuf* **CAG1** |  |  |  |  |
|  | *Glaesserella parasuis* | -0.26 | 16S | CAG1 |  | *Gammaproteobacteria* Class | -0.30 | *tuf* | CAG2 |
|  | *Phascolarctobacterium succinatutens* | -0.21 | 16S | - |  | *Lactobacillales* Order | -0.28 | *tuf* | CAG2 |
|  | *Lactobacillus reuteri* | -0.21 | 16S | CAG4 |  | *Moraxella* Genus | -0.28 | *tuf* | CAG2 |
|  | *Mannheimia caviae* | -0.2 | 16S | CAG1 |  | *Moraxella osloensis* | -0.26 | *tuf* | CAG2 |

## Discussion

In piglets the development of the nasal microbiome has implications for health and disease, but longitudinal development is understudied, and most microbiome studies lack taxonomic resolution We obtained longitudinal data of the porcine nasal microbiota from birth to day 70 of life, from three farms across three different countries, using a combination of 16S rRNA and *tuf* gene amplicon sequencing achieving phylogenetic resolution up to species level. Using *tuf* sequencing the species resolution of *Firmicutes* (34, 35) in the piglet nasal microbiome improved. This allowed us to describe CAGs of bacterial species that are commonly found together and consistently present in piglets in the studied countries.

### Piglet age and the farm shape the nasal microbiota

Time (age of the piglet) explained 19.4% of the variation in nasal microbiome composition between samples. The microbiome of pigs changed rapidly over time, as previously observed in human (36) and animal (37–39) studies, and a more stable microbiome has been observed in adulthood (36, 40). Weaning is an impactful life event for piglets shown in this study by stabilisation of the PNM composition between weaning T=27 and two weeks post weaning (T=41). This post-weaning stabilisation has previously been described (41) and suggests maturation from juvenile to a more “adult” microbiome in response to the changing factors around weaning.

After age, the farm it was raised on had the largest effect on beta diversity, followed by its sow (litter), and the country of origin (Table 2). A farm effect (9) and a litter effect on the piglet-tonsil microbiota (42) have previously been described. Nonetheless, remarkable congruence in the developing microbiome was observed for all piglets across all countries. Homogeneity in management and genetics in commercial pig farming might help explain this, despite pig lineage and origin differences between farms in this study. To determine drivers behind the difference between farms, countries, and/or litters that effect nasal microbiome development, a study design with more farms and farm data would be needed.

### The vaginal and faecal microbiome ‘seeds’ the nasal microbiome

Previously, we observed that nasal samples taken from piglets directly after birth contained many species associated with the gastrointestinal tract (7). Inception of the piglets’ microbiome development through birth canal passage and first contact with the environment (including faeces) is supported by the altered development of the (gut) microbiome in C-section derived pigs (43).

The sow microbiome and the sow’s parity influence the piglet tonsillar microbiome directly post-partum. As seen in the vaginal microbiota of high parity sows consisting of taxa as *Pasteurellaceae* family, and the *Aerococcus, Clostridium,* and *Escherichia* genera. These taxa were subsequently more present in their piglets’ tonsil directly *post-partum* (31). Similarly, in our samples shortly after birth we detected members of the genera *Clostridium* and *Escherichia* (Figure 2). Clostridium species clustered into 16S-CAG2 and 16S-CAG3 (16S-CAG3 almost completely consisted of *Clostridiaceae*). Both CAGs were detected mostly in early life (CAG3 has an additional peak around week two). The abundance of these taxa and the higher Shannon alpha diversity (Figure 1) indicate nasal colonization by the generally rich vaginal and faecal microbiota. A recent study shows that gut associated microbes are present nasally and actively living there (44). These observations add further evidence that natural birth seeds the early colonizers of the piglet’s upper respiratory tract.

### Sow and farrowing pen conditions shape the starting microbiome

The samples from the first week after birth excluding the first timepoints (T=2-6) likely consist of a microbiome that is driven by maternal or farrowing pen influences as the samples of these timepoints cluster together horizontally. Colostrum and the high piglet snout to sow skin/teat contact may shape the microbiota as has been suggested by Obregon-Gutierrez and collaborators. Specifically, sow-skin contact was associated with presence of the genera *Rothia*, *Moraxella*, and *Enhydrobacter* (45). *Enhydrobacter*, a member of the *Moraxallaceae* family, was not observed in our study. However, we did observe high abundances of *Rothia* but was less abundant than *Moraxella* and *Mannheimia* species mainly present during the first week of life. This is supported by our previous finding that *Rothia* was present at almost 50% relative abundance during the first week (7). *M. boevrei* was often the most prevalent species during this period, succeeded later in life by other *Moraxella* species, a finding consistent with other studies (10, 12, 46). Interestingly, *Moraxella* has been observed as a common nasopharyngeal inhabitant in humans and animals and has been associated with opportunistic airway infections (47–50), but also with breastfeeding, and microbiome stability (40).

### Abundant taxa detected in the nasal microbiota match previous studies

In alignment with our data (Figure S.4), Correa-Fiz *et al*. (2016) identified genera such as *Moraxella, Haemophilus, Enhydrobacter, Klebsiella, Oscillospira, Streptococcus, Lactobacillus, Weeksella, Prevotella, Bacteriodes* as key components of the core nasal microbiome at 3-4 weeks of life (10). The discrepancy between *Haemophilus* and *Glaesserella* can be explained by a name change of *H. parasuis* to *G. parasuis*. The most abundant genera in their 2019 study (46), again correlate with our current findings. The porcine nasal microbiome samples studied by Strube *et al.* (2018) (51) had higher *Streptococcus*, *Rothia, Moraxella,* and *Globicatella* and lower *Facklamia* and *Aerococcus* abundances. We observed these genera, except *Facklamia* and *Globicatella,* as the most abundant taxa. The most abundant genera in our study, are in direct concordance with the genera present in the pig upper respiratory microbiome, as found at three timepoints by *Rampelotto et al*. (2022) at the end of weaning, at the end of the nursery phase (T=71), and at finishing (T=373) (12).

### Co-abundance analysis identifies two major CAGs in competition

In both 16S rRNA and *tuf* datasets co-abundant species grouped across countries. Previously, we described CAGs from 16S rRNA and *tuf* datasets in one Dutch farm with temporal trends and associations with LA-MRSA (7). Here we describe seven 16S-CAGs across multiple farms and countries. There was overlap between 16S-CAG6 and 16S-CAG8 of the previous study, and 16S-CAG1 and 16S-CAG3 of the current study. For example, in previous 16S-CAG8 and current 16S-CAG3 we observe *Clostridium* species being present two weeks pre-weaning.

The interaction network depicts that the members of the two largest 16S-CAGs are negatively associated with each other (Figure 5A). The same holds true for the two *tuf*-CAGs we defined (Figure 5B). These negative associations have a temporal component. Members of 16S-CAG1 and *tuf*-CAG2 are most present in the later time points, indicating that these CAGs are part of the maturated microbiome. We argue that there is a group of genera colonizing the nasal microbiome at birth and early in life (for example the species in 16S-CAG2). This group is later succeeded by other taxa (the species in 16S-CAG1) potentially due to microbe-competition, changing maternal contact, piglet development or by altered piglet behaviour and changes in its diet (e.g. weaning or early life solid feed introduction).

### Taxa antagonistic to respiratory pathogens

We observed widespread presence of opportunistic pathogens, with low relative abundances, and observed farm specific pathogen trends. Several taxa were negatively associated with pathogens (Table 4). As example, some farms showed *M. hyorhinis* in co-occurrence with *P. multocida* after week two. This is in concordance with clinical signs of *M. hyorhinis* infections arising around week 3-10 in affected pigs (52) and matches previous descriptions of secondary infections of *M. hyorhinis*, *P. multocida* and *S. suis* after viral infection as part of the PRDC complex (53).

However, piglets in our study did not show clinical symptoms. This absence of symptoms is consistent with studies that detected bacteria commonly associated with disease without causing disease (19, 54). Therefore, these species could be considered as opportunistic pathogens or pathobionts (55).

Before using the current findings to develop competitive exclusion against respiratory pathogens, additional experimental evidence is warranted. The amplicon sequencing used in the current study resulted in compositional data which is not ideal for quantitative analysis (56). Absence of taxa in the sequence data does not necessarily indicate a true absence of the bacterium (7). Furthermore, methodological differences can effect on the outcome of a sequencing effort and may cause differences between studies (57). Quantitative and sensitive methods such as qPCR are needed to confirm negative correlations between potential probiotic taxa and opportunistic pathogens. Additionally, interactions identified in our *in silico* analyses, should be tested by isolating identified strains and testing them in *in vitro* and *in vivo* models.

## Conclusion

This study extends the understanding of the longitudinal development of the piglet nasal microbiome. The nasal microbiota of 54 piglets between birth and day 70 of life showed comparable developmental trends in three geographically distinct regions, different farms, and litters. We conclude that the development of the piglet nasal microbiome is strongly influenced by age of the pig regardless of the rearing location. CAGs of bacterial species, with distinct temporal trends in all three countries, showed interactions between commensal nasal species and respiratory pathogens. These data can be used to inform and identify targeted probiotic strains/probiotics to control opportunistic bacterial airway pathogens. Such a pathogen reduction strategy would be valuable in the fight against AMR.

## Material & Methods

### Farm selection and animal management

Nine conventional farms were selected in three European countries (Ireland (n=3), Germany (n=3), and the Netherlands (n=3)). Farmers gave informed consent to participate in the study and all data were anonymised before analysis. At birth, piglets from four litters on each of the three farms in the three countries were selected for nasal swabbing. Only piglets from sows which had not been treated with antimicrobials were selected, and piglets from litters where antimicrobials were administered during the study period were excluded. In the Netherlands, sampling protocols were reviewed and approved by the Animal Ethical Committee of Utrecht University, the Netherlands (Declaration no. 111218). In Ireland, Ethical approval for the study was granted by the Teagasc Animal Ethics Committee (approval no. TAEC207-2018) and the project was authorised by the Irish Health Products Regulatory Authority (project authorisation no. AE19132/P094). The experiment was conducted in accordance with EU standards (European Directive 2010/63/EU), the legislation for commercial pig production set out in the European communities (welfare of farmed animals), and in Irish legislation (SI no. 311/2010). Farm specific management was maintained, and all materials were collected by trained veterinarians/animal scientists.

### Nasal swab sampling

Piglets were nasal swabbed at 0, 1, 2, 3, 4, 5, 6, 13, 20, 27, 34, 41, 48, 55, 62, and 69 days after birth. Nasal swabs were taken equally from 54 piglets born from 27 sows across nine farms over three countries (N = 864). For samples taken in week one, aluminium-wire swabs with a rayon tip (Copan, 160C) were used. Swabs from later timepoints (T=14-69) were taken using plastic swabs with a rayon tip (Copan, 155C). The swabs were kept refrigerated and were processed within 24 hours post sampling. The swab tips were cut into 2 mL Eppendorf tubes and submerged in 600 µL lysis buffer BL (LGC genomics, Berlin Germany) and stored at −20^0^C until processing.

### DNA extraction

DNA from nasal swabs was extracted, from piglets with a completed timeseries (n=16) (for Dutch farm NLD3 (n=9)) resulting into a total of 813 samples, using a modified LGC mag kit protocol (LGC genomics, Berlin, Germany) adapted from Wylie *et al.* (58). The samples were homogenised and lysed by beat beating with 0.2 mm zirconia beads and 500 µL molecular grade TE saturated phenol (ThermoFischer, Bleiswijk, the Netherlands). After spinning the aqueous phase was transferred to 1000 µL binding buffer and 10 µL magnetic beads in a round bottom molecular grade 2.2 mL 96 deep wells plate (VWR international, Amsterdam, the Netherlands). The binding and washing steps were performed in 96 well plates according to the manufacturer’s specification, and the eluted DNA was stored at −20^0^C.

### Quantification of 16S rRNA by real-time PCR to normalise sequencing input

Sample bacterial DNA was estimated using a 16S rRNA qPCR to normalise the sequencing library preparation. Reactions were performed on the LightCycler 480 platform (Roche Diagnostics, Almere, The Netherlands). Reaction mixtures consisted of 1 µL nasal swab DNA, 7 µL molecular grade water, 1 µl primer 355F (5’ ACTCCTACGGGAGGCAGC 3’) at 10 µM, 1 µL primer 556R (5’ CTTTACGCCCARTRAWTCCG 3’) at 10 µM and 10 µL SYBER® Green Master Mix (Bio-Rad, Veenendaal, The Netherlands). The DNA extraction estimated an average rRNA yield corresponding to 5×10^8 bacterial cells/mL.

### 16S rRNA and *tuf* gene sequencing

Amplicon libraries targeting the V3-V4 region of the 16S rRNA gene was amplified using the primers 341F (5′-TCGTCGGCAGCGTCAGATGTGTATAAGAGACAGC CTACGGGNGGCWGCAG-3′) and 805R (5′-GTCTCGTGGGCTCGGAGATGTGTATAAGA GACAGGACTACHVGGGTATCTAATCC-3′) (Eurofins Genomics, Germany), and the *tuf* gene was amplified using the primers tuf-F (5′-GCCAGTTGAGGACGTATTCT-3′) and tuf-R (5′-CCATTTCAGTACCTTCTGGTAA-3′). Both primer sets were used at a final concentration of 0.2 µM with Phusion High-Fidelity DNA polymerase (Thermo Scientific, USA). According to Illumina (San Diego, CA, USA) recommendations, the following PCR conditions were used: an initial denaturation step at 98 °C for 30 s, followed by 25 cycles of denaturation at 98 °C for 10 s, annealing at 55 °C for 15 s, extension at 72 °C for 20 s, and a final extension step at 72 °C for 5 min, followed by cooling to 4 °C. The amplicons were visualized on a 1% agarose gel to confirm amplicon lengths of approximately 550 bp for the 16S rRNA gene and 400 bp for the *tuf* gene. The amplicons were purified using Agencourt AMPure XP magnetic beads (Beckman-Coulter, USA) and eluted in 50 µL EB Buffer (Qiagen), after which Illumina barcode sequences were added using the Nextera XT v2 Index primer kit. The index PCR conditions were as follows: 98 °C for 30 s, followed by 8 cycles of denaturation at 98 °C for 10 s, 55 °C for 15 s, 72 °C for 20 s, and a final 72 °C for 5 min, followed by cooling to 4 °C. Another round of purification with Agencourt AMPure XP magnetic beads was performed and the final amplicons with indices, were eluted in 28 µL EB Buffer (Qiagen). Sample concentration was measured using the Qubit high-sensitivity double-stranded DNA assay kit (Thermo Scientific) on a Qubit™ 3 Fluorometer. For library pooling, 30 ng DNA from each sample was combined to create a randomized pooled library, that were sequenced at the Teagasc NGS sequencing facility (Teagasc Moorepark, Fermoy, Co. Cork, Ireland) on an Illumina MiSeq, generating 2 × 300 bp paired-end reads.

### Microbiome pre-processing

The check of read quality and the pipeline to infer Ribosomal Sequence variants (RSVs) using DADA2 v1.20 was performed as described by Patel *et al.*, 2021 (7) with slight modifications considering the following parameters: truncLen=c(220,220), trimLeft=c(17,21), maxEE=c(2,2), truncQ=c(2,2), maxN=0, rm.phix=TRUE for 16S rRNA and truncLen=c(240,200), trimLeft=c(20,22), maxEE=c(2,2), truncQ=c(2,2), maxN=0, rm.phix=TRUE for the *tuf* gene. In the taxonomy assignment step the bootstrap confidence threshold was mutated to 80%.

### Microbiome analysis

Datasets were formatted into two phyloseq-objects (*tuf* and 16S rRNA) for diversity analyses, visualisation, and statistics. RStudio for windows (2023.03.1+446 “Cherry Blossom” Release) with R version 4.1.3 (2022-03-10) was used to perform all analyses. Analyses in R were performed using the “microViz” (0.10.6) (59), “microbiome” (1.16.0), “tidyverse” (1.3.2), “CoDaSeq” (0.99.6), “phyloseqCompanion” (1.0), “vegan” (2.6-4), and “phyloseq” (1.38.0) packages. The full R analysis, including all loaded libraries (utilitarian libraries), is appended as a Quarto-document (.qmd).

### Alpha Diversity calculations

Alpha diversity measures were estimated and visualised using the “phyloseq” R-package. To correct for varying sequence depth all samples were rarefied to 5034 reads for the 16S rRNA dataset (minimum read count of the 16S rRNA dataset) and 6091 reads for the *tuf* dataset (minimum read count of the *tuf* dataset). Shannon alpha diversity was visualised using the ‘plot_richness()’ function. T-tests to compare timepoint alpha diversity were performed using the base R Welch two sample T-test.

### Beta diversity calculations

Aitchison distances were calculated from centered log-ratio (CLR) transformed, unfiltered, unrarefied datasets which had zeroes imputed with half the minimum observed value for each taxon at species level, using the ‘tax_transform()’ function. When taxonomy was not assignable to species level, the microViz “tax_fix()” function was used to impute the highest possible taxonomic resolution (E.G. *Staphylococcus* species). Samples were ordinated using the ‘ord_calc()’ function using method “PCA”. The ordinations were visualised using the ‘ord_plot()’ function. All three functions are from the “microViz” R-package.

### PERMANOVA analysis on beta diversity

PERMANOVA analysis was performed on the ordinations mentioned above using the ‘adonis2()’ function from the “Vegan” R-package. Using the model formula: “Country/Farm/Identifier_SOW/Identifier_pig + Time + Sex” to consider the nested study design. We used 5000 permutations.

### Composition bar plots and heatmaps

Composition bar plots were generated by merging relative abundance data from all farms by timepoint for each country. Unrarefied datasets were used to generate the composition bar plots. Bar plots were generated using the ‘comp_barplot()’ function and heatmaps were generated using the ‘ cor_heatmap()’ function both from the “microViz” R-package, setting tax_level to “species”, and n_taxa to 25. Figures were optimised using functions from the “ggplot2” R package.

### SparCC correlation

ASVs were agglomerated to species level to generalize species level effects. Positive and negative associations based on the abundances of species with at least 50 reads in at least 5% of the samples in the 16S rRNA and *tuf* dataset, were inferred using SparCC (60). From each separate country (anti)correlations with an r >0.2 or r <-0.2 were selected. Species displaying significant correlation with an average correlation of r > 0.2 over all countries where the interaction was found, were clustered into co-abundance groups (CAGs) using MCL (61) using default settings.

## Acknowledgements

We want to thank Marian Broekhuizen-Stins, Gerard van Eijden, Arjen Timmerman, and Heleen Zweerus, for their contributions to the sampling, and sample-processing effort.

## Funding

This research was funded by The Irish Health Research Board (ExcludeMRSA; grant number JPI-AMR-2017-1-A) and in part by Science Foundation Ireland (grant numbers SFI/12/RC/2273_P2 and 17/CDA/4765).

The Dutch part was supported by ZonMw, and JPIAMR (JPIAMR-2017-1-B grant number 50-52900-98-043).

## Availability of data and materials

Scripts and Phyloseq objects are available through ZENODO Reads are available under accession number: PRJEB71383

## Authors’ contributions

AZ, BD, JAW, PL, CE and MJC designed the study. DCP, PL, AAV, JAW and CE collected the pig samples. JE, CH and AAV carried out the amplicon sequencing. SP, AAV, AZ, RL, SP performed the data analysis. AAV, MJC, BD, AZ, JAW, PL, and RL participated in the interpretation of the results. AAV wrote the manuscript with input from all other authors. All authors read and approved the final manuscript.

## Ethics approval and consent to participate

See statement in Material and Methods

## Consent for publication

Informed consent by the farmers was obtained.

## Competing interests

None stated.

## Appendixes

**S. 1A.**
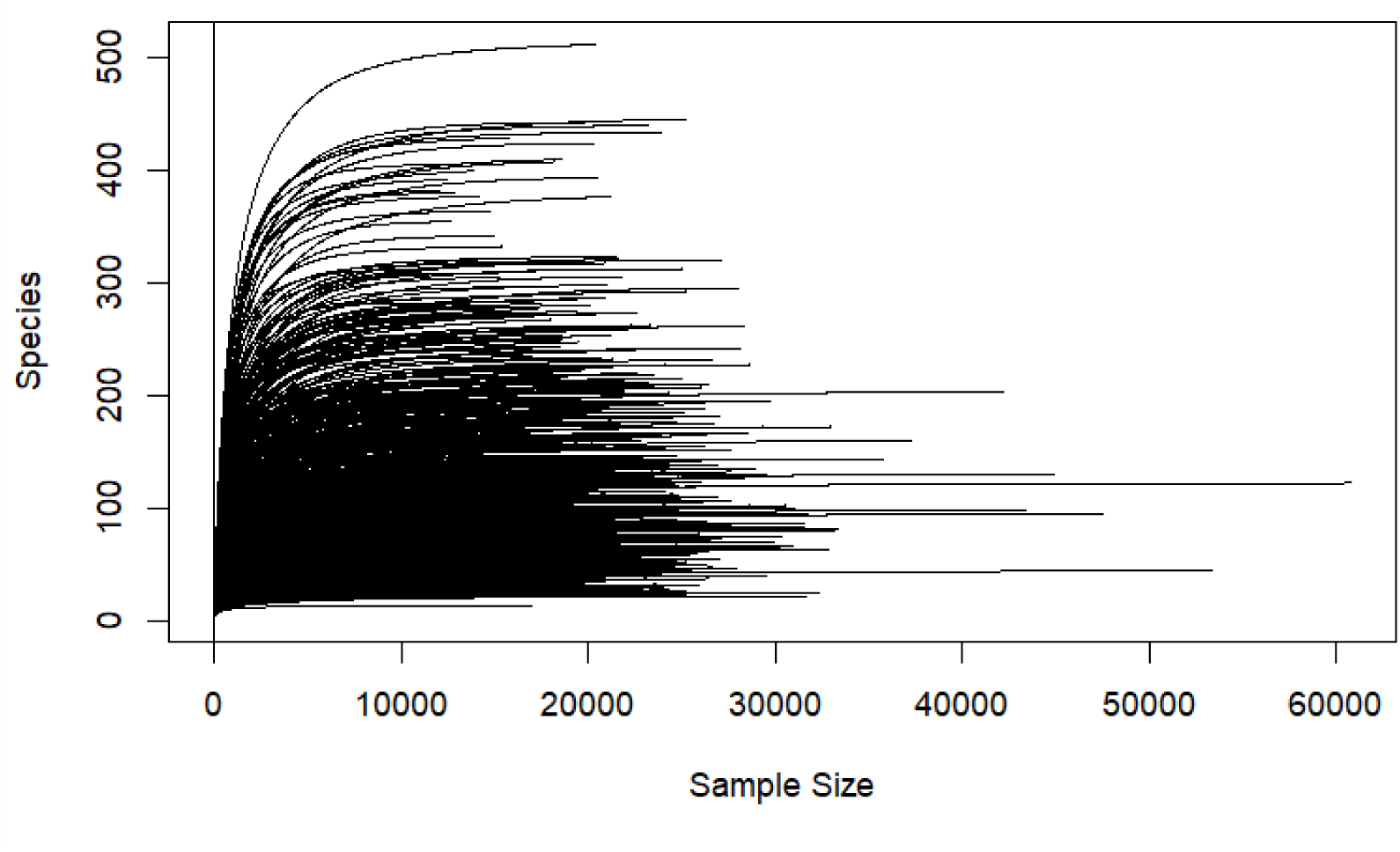
Rarefaction curve of all piglets 16S rRNA samples pre rarefaction. This indicates that sufficient read depth has been achieved to represent the RSVs in the sequenced 16S rRNA gene.

**S. 1B.**
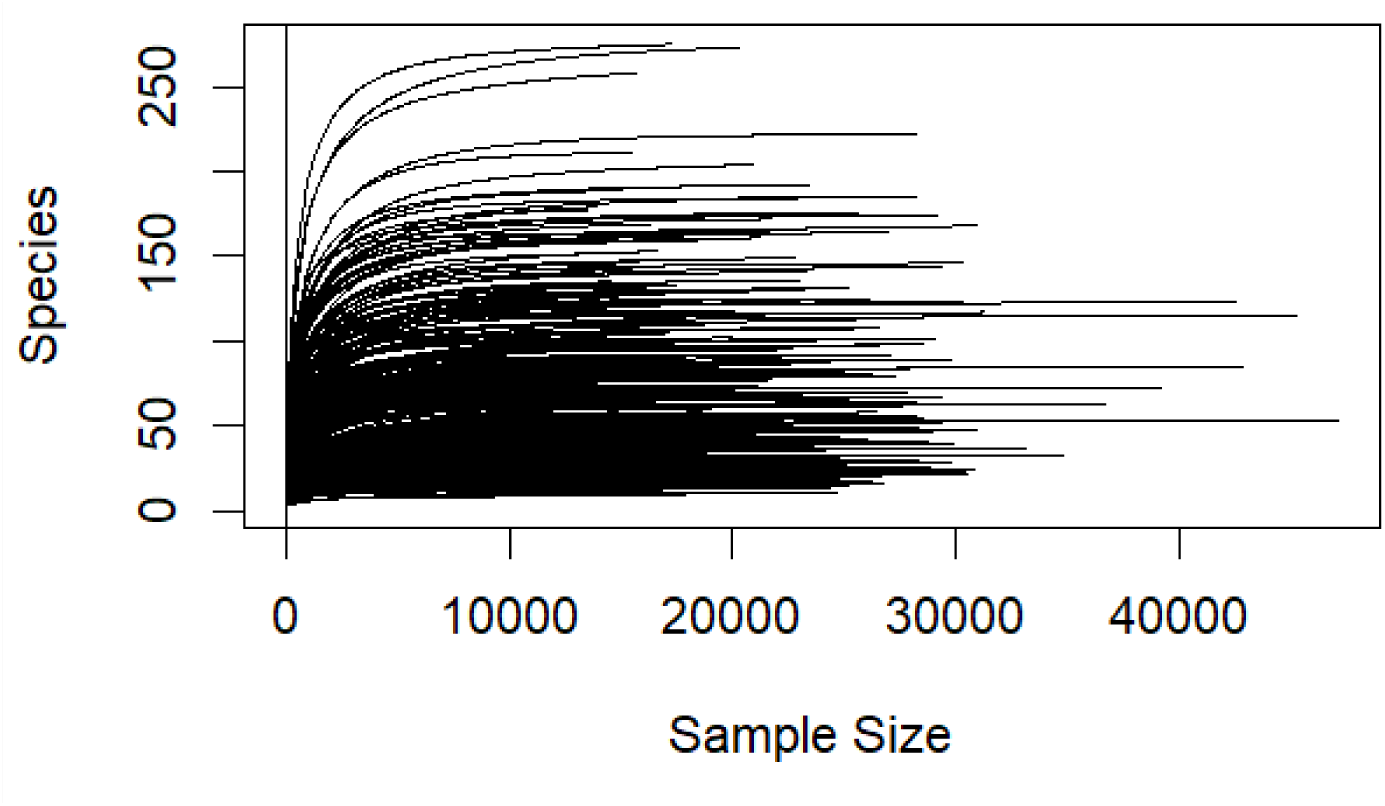
Rarefaction curve of all piglets *tuf* samples pre rarefaction. This indicates that sufficient read depth has been achieved to represent the TSV’s in the sequenced *tuf* genes.

**S. 2.**
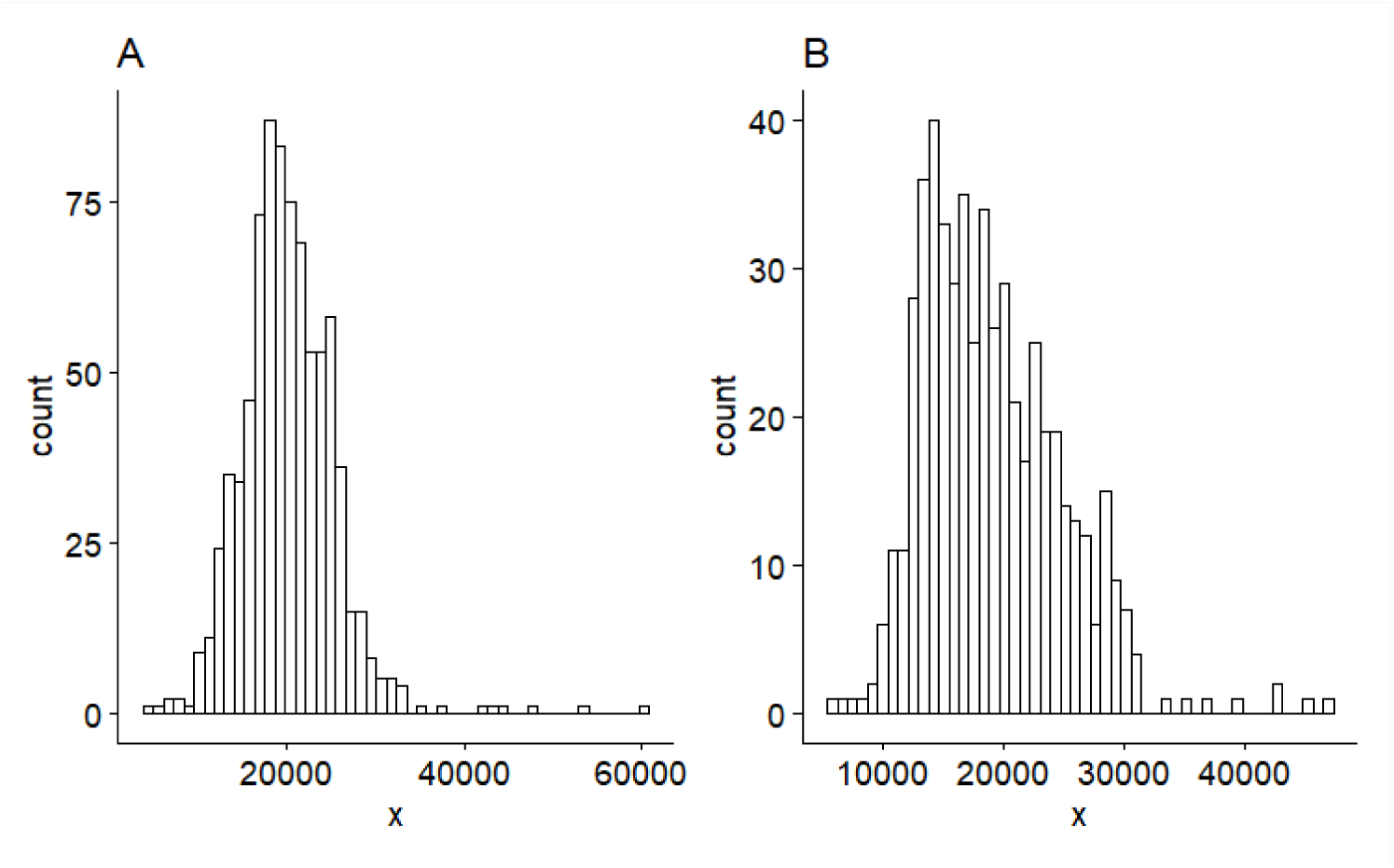
**A** Distribution of read depth 16S rRNA gene sequencing **B** Distribution of read depth *tuf* sequencing

**S. 3.**
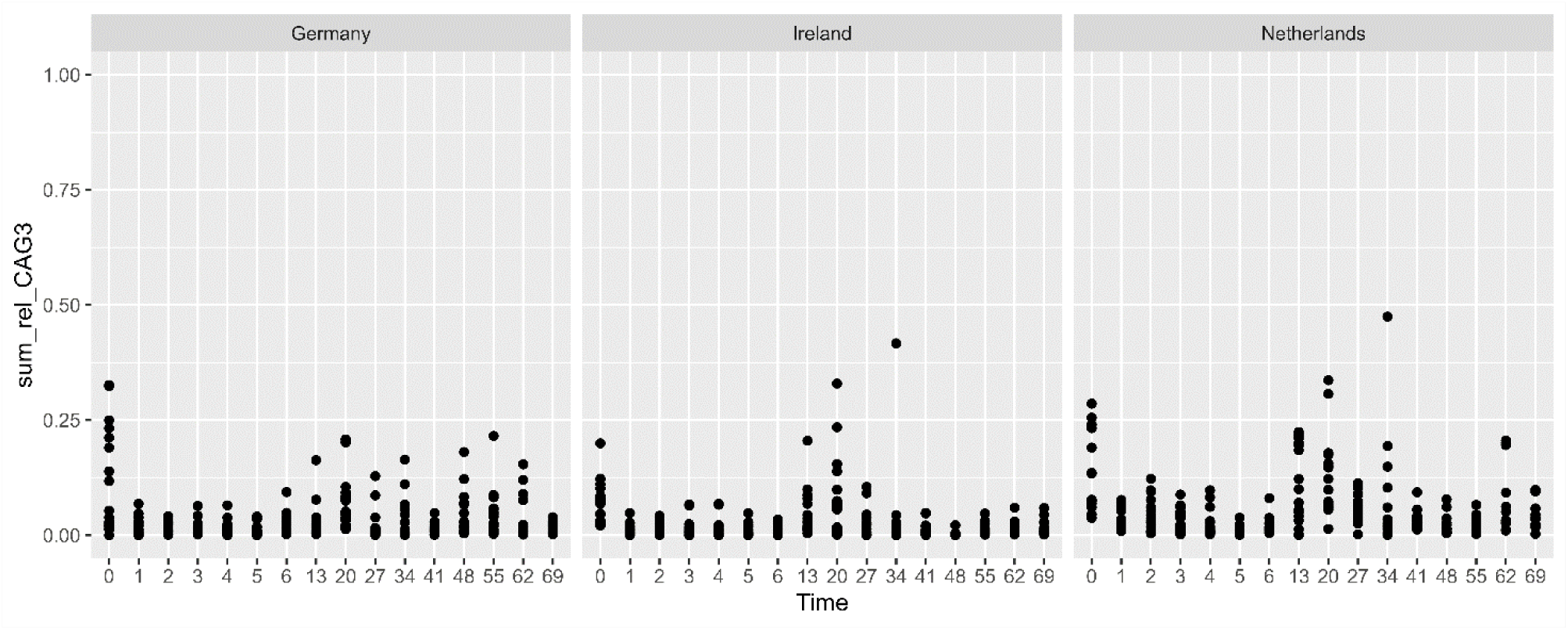
Dotplot displaying relative abundances of 16S-CAG3 per sample. Relative abundance of 16S-CAG3 spikes around weaning (T27).

**S.4.**
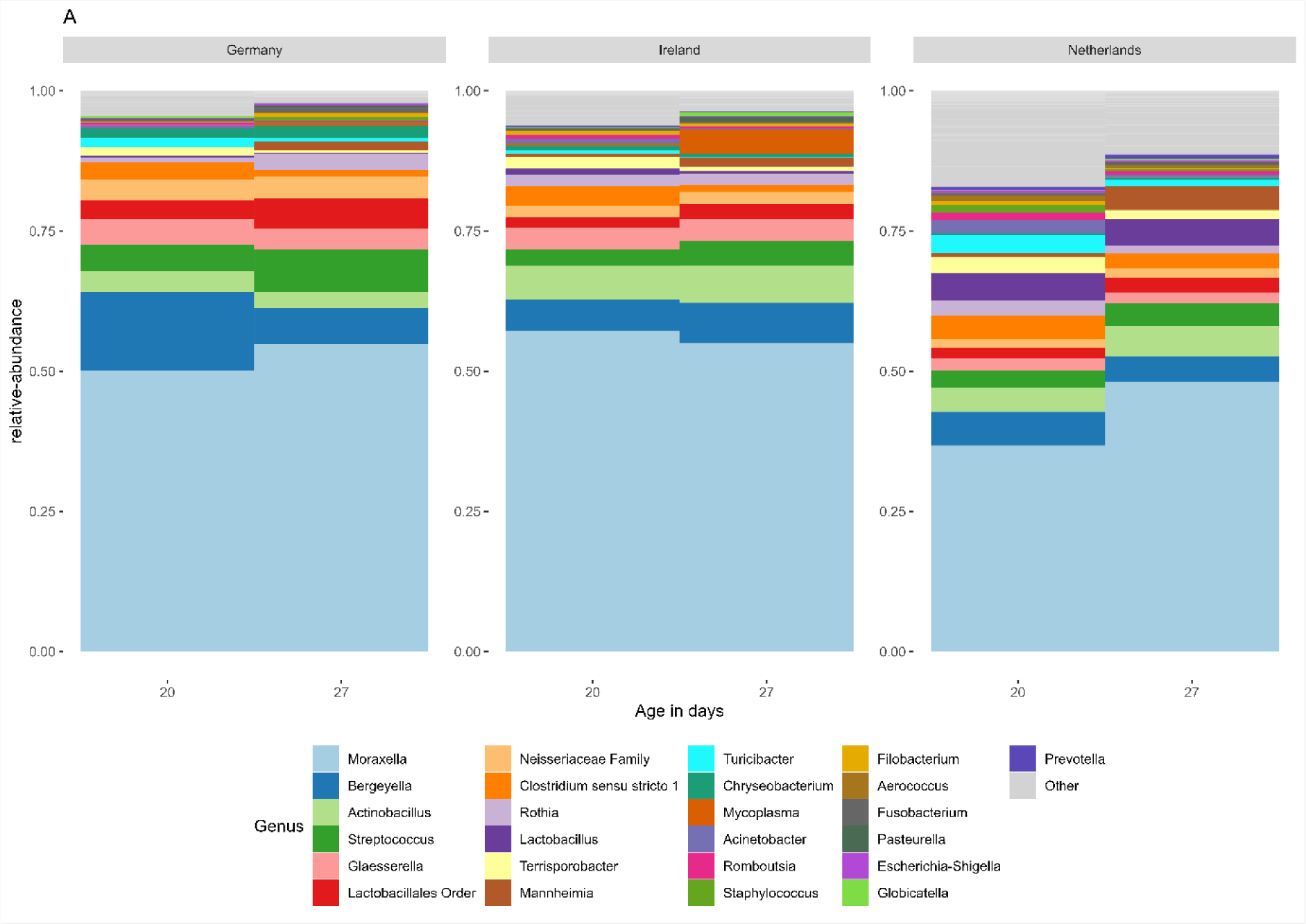
Barplots displaying relative abundances of the top 25 taxa at genus level for timepoint week three and week four, compressed by country.

**S.5.**
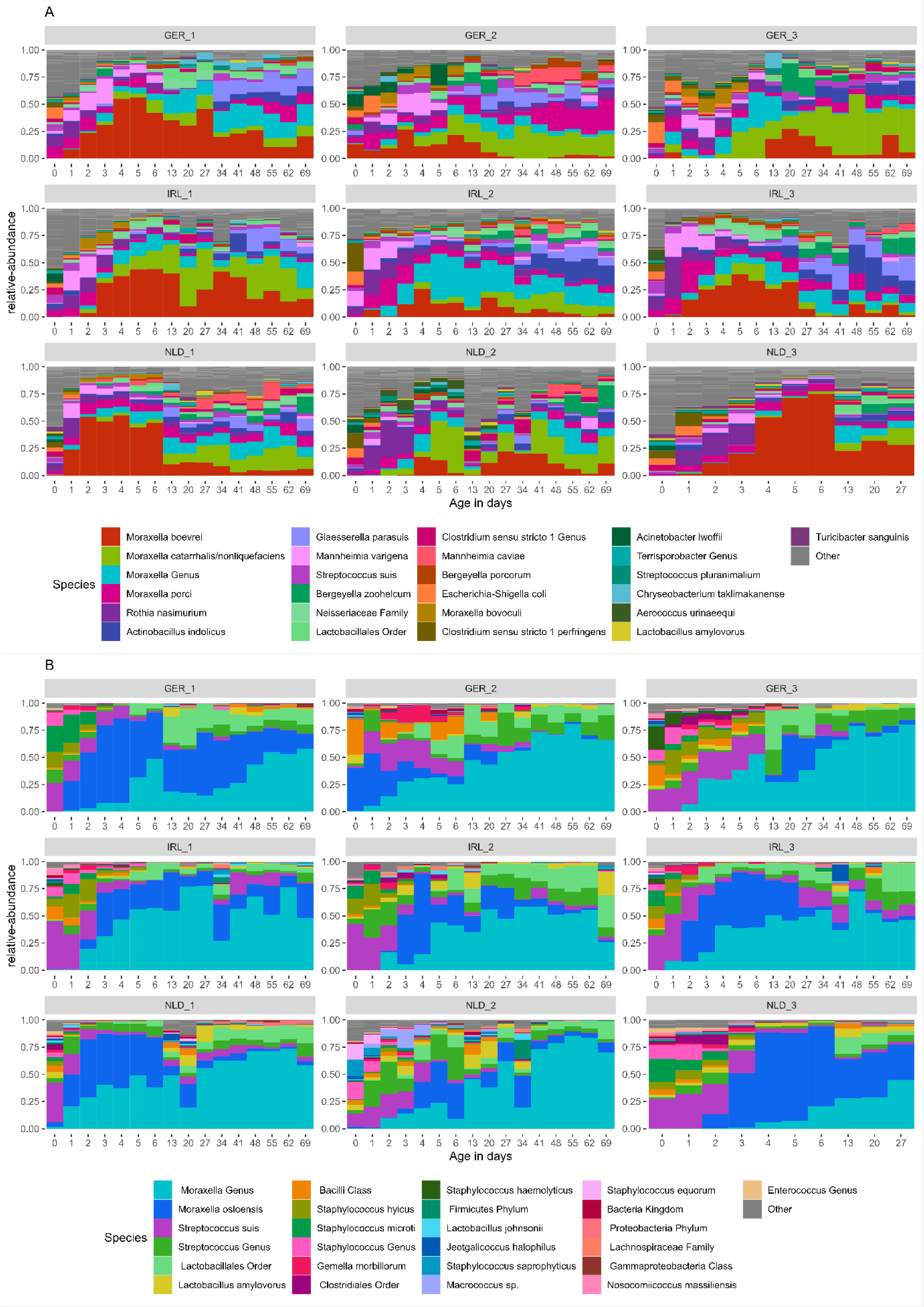
**A** Top 25 species from the 16S rRNA dataset of all samples compressed into a one bar per timepoint per farm. **B** Top 25 species from the *tuf* dataset of all samples compressed into a one bar per timepoint per farm.

**S.6.**
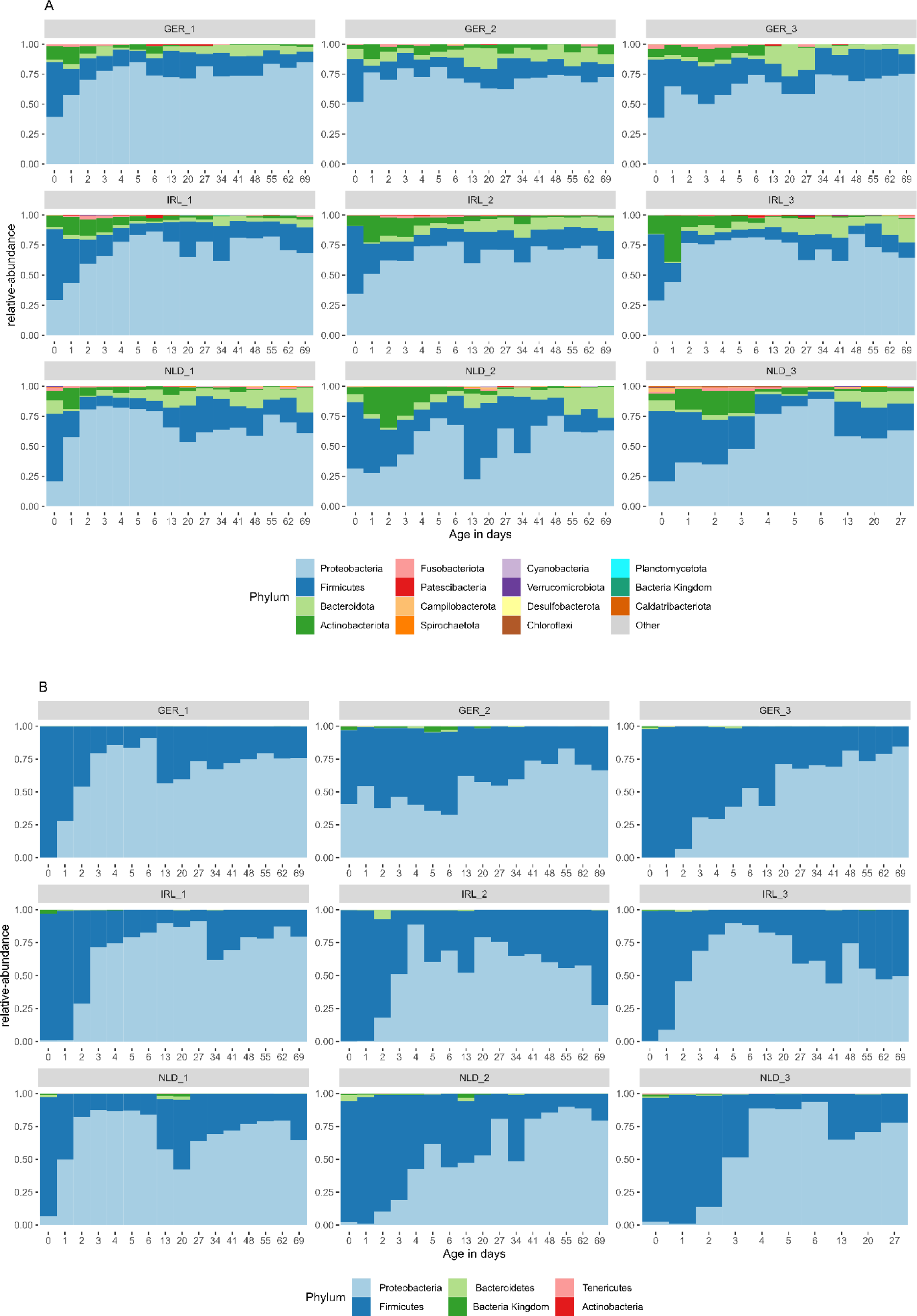
**A** 16S rRNA Phyla per farm in time **B** *tuf* Phyla per farm in time.

**S.7.**
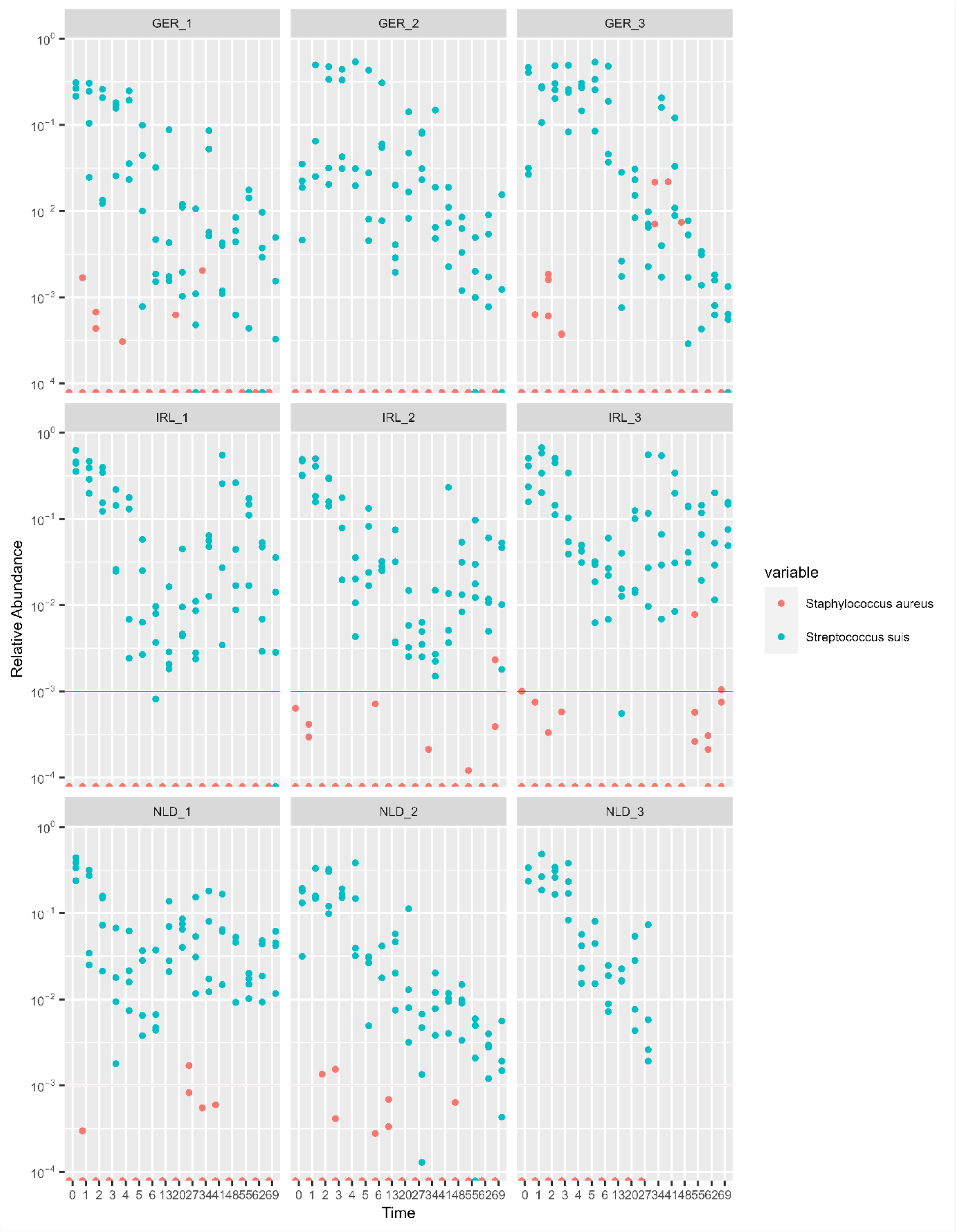
*tuf* pathogens relative abundances per sample in time.

**S. Table 1.**
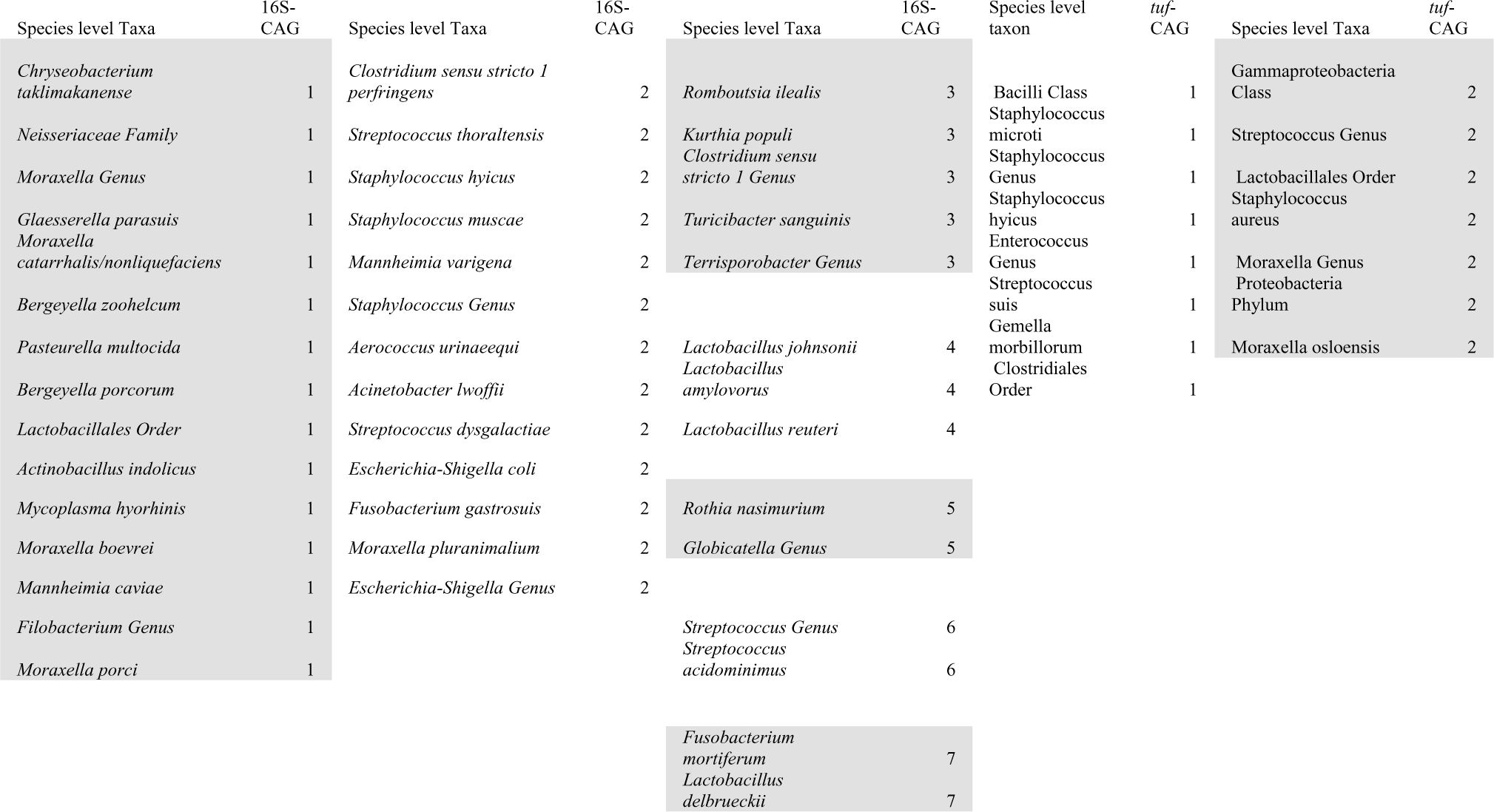
Co-abundance groups (CAGs) in the 16S rRNA and *tuf* dataset with an average co-correlation of r>0.2 over all three countries.

## References

1. Opriessnig T, Giménez-Lirola LG, Halbur PG. 2011. Polymicrobial respiratory disease in pigs. Anim Health Res Rev 12:133–148.

2. Ruggeri J, Salogni C, Giovannini S, Vitale N, Boniotti MB, Corradi A, Pozzi P, Pasquali P, Alborali GL. 2020. Association Between Infectious Agents and Lesions in Post-Weaned Piglets and Fattening Heavy Pigs With Porcine Respiratory Disease Complex (PRDC). Front Vet Sci 7:636.

3. Amat S, Alexander TW, Holman DB, Schwinghamer T, Timsit E. 2020. Intranasal Bacterial Therapeutics Reduce Colonization by the Respiratory Pathogen Mannheimia haemolytica in Dairy Calves. mSystems 5:1–19.

4. Piewngam P, Zheng Y, Nguyen TH, Dickey SW, Joo HS, Villaruz AE, Glose KA, Fisher EL, Hunt RL, Li B, Chiou J, Pharkjaksu S, Khongthong S, Cheung GYC, Kiratisin P, Otto M. 2018. Pathogen elimination by probiotic Bacillus via signalling interference. Nat 2018 5627728 562:532–537.

5. Piewngam P, Khongthong S, Roekngam N, Theapparat Y, Sunpaweravong S, Faroongsarng D, Otto M. 2023. Probiotic for pathogen-specific Staphylococcus aureus decolonisation in Thailand: a phase 2, double-blind, randomised, placebo-controlled trial. The Lancet Microbe 4:e75–e83.

6. Pirolo M, Espinosa-Gongora C, Bogaert D, Guardabassi L. 2021. The porcine respiratory microbiome: recent insights and future challenges. Anim Microbiome 3:1–13.

7. Patel S, Vlasblom AA, Verstappen KM, Zomer AL, Fluit AC, Rogers MRC, Wagenaar JA, Claesson MJ, Duim B. 2021. Differential Analysis of Longitudinal Methicillin-Resistant Staphylococcus aureus Colonization in Relation to Microbial Shifts in the Nasal Microbiome of Neonatal Piglets. mSystems 6:e0015221.

8. Niederwerder MC. 2017. Role of the microbiome in swine respiratory disease. Vet Microbiol 209:97–106.

9. Arruda AG, Deblais L, Hale VL, Madden C, Pairis-Garcia M, Srivastava V, Kathayat D, Kumar A, Rajashekara G. 2021. A cross-sectional study of the nasal and fecal microbiota of sows from different health status within six commercial swine farms. PeerJ 9:e12120.

10. Correa-Fiz F, Fraile L, Aragon V. 2016. Piglet nasal microbiota at weaning may influence the development of Glässer’s disease during the rearing period. BMC Genomics 17:1–14.

11. Mahmmod YS, Correa-Fiz F, Aragon V. 2020. Variations in association of nasal microbiota with virulent and non-virulent strains of Glaesserella (Haemophilus) parasuis in weaning piglets. Vet Res 51:1–13.

12. Rampelotto PH, dos Santos ACR, Muterle Varela AP, Takeuti KL, Loiko MR, Mayer FQ, Roehe PM. 2022. Comparative Analysis of the Upper Respiratory Bacterial Communities of Pigs with or without Respiratory Clinical Signs: From Weaning to Finishing Phase. Biology (Basel) 11:1111.

13. Li Z, Wang X, Di D, Pan R, Gao Y, Xiao C, Li B, Wei J, Liu K, Qiu Y, Ma Z. 2021. Comparative analysis of the pulmonary microbiome in healthy and diseased pigs. Mol Genet Genomics 296:21–31.

14. Wang Q, Cai R, Huang A, Wang X, Qu W, Shi L, Li C, Yan H. 2018. Comparison of Oropharyngeal Microbiota in Healthy Piglets and Piglets With Respiratory Disease. Front Microbiol 9:3218.

15. Niederwerder MC, Jaing CJ, Thissen JB, Cino-Ozuna AG, McLoughlin KS, Rowland RRR. 2016. Microbiome associations in pigs with the best and worst clinical outcomes following co-infection with porcine reproductive and respiratory syndrome virus (PRRSV) and porcine circovirus type 2 (PCV2). Vet Microbiol 188:1–11.

16. Weese JS, Slifierz M, Jalali M, Friendship R. 2014. Evaluation of the nasal microbiota in slaughter-age pigs and the impact on nasal methicillin-resistant Staphylococcus aureus (MRSA) carriage. BMC Vet Res 10:1–10.

17. Espinosa-Gongora C, Larsen N, Schønning K, Fredholm M, Guardabassi L. 2016. Differential analysis of the nasal microbiome of pig carriers or non-carriers of staphylococcus aureus. PLoS One 11:1–13.

18. Wang T, He Q, Yao W, Shao Y, Li J, Huang F. 2019. The Variation of Nasal Microbiota Caused by Low Levels of Gaseous Ammonia Exposure in Growing Pigs. Front Microbiol 10:1083.

19. Fablet C, Marois-Créhan C, Simon G, Grasland B, Jestin A, Kobisch M, Madec F, Rose N. 2012. Infectious agents associated with respiratory diseases in 125 farrow-to-finish pig herds: A cross-sectional study. Vet Microbiol 157:152–163.

20. Saade G, Deblanc C, Bougon J, Marois-Créhan C, Fablet C, Auray G, Belloc C, Leblanc-Maridor M, Gagnon CA, Zhu J, Gottschalk M, Summerfield A, Simon G, Bertho N, Meurens F. 2020. Coinfections and their molecular consequences in the porcine respiratory tract. Vet Res 2020 511 51:1–19.

21. Gamage D, Sulochana R, Alaa K□, Fakher A, Matheus De Oliveira Costa □. 2022 Actinobacillus suis isolated from diseased pigs are phylogenetically related but harbour different number of toxin gene copies in their genomes. Vet Rec Open 9:e45.

22. Jarosz ŁS, Gradzki Z, Kalinowski M. 2014. Trueperella pyogenes infections in swine: clinical course and pathology. Pol J Vet Sci 17:395–404.

23. Sarli G, D’Annunzio G, Gobbo F, Benazzi C, Ostanello F. 2021. The Role of Pathology in the Diagnosis of Swine Respiratory Disease. Vet Sci 8.

24. Kehrenberg C, Salmon SA, Watts JL, Schwarz S. 2001. Tetracycline resistance genes in isolates of Pasteurella multocida, Mannheimia haemolytica, Mannheimia glucosida and Mannheimia varigena from bovine and swine respiratory disease: intergeneric spread of the tet(H) plasmid pMHT1. J Antimicrob Chemother 48:631–640.

25. Clavijo MJ, Davies P, Morrison R, Bruner L, Olson S, Rosey E, Rovira A. 2019. Temporal patterns of colonization and infection with Mycoplasma hyorhinis in two swine production systems in the USA. Vet Microbiol 234:110–118.

26. Jirawattanapong P, Stockhofe-Zurwieden N, van Leengoed L, Wisselink H, Raymakers R, Cruijsen T, van der Peet-Schwering C, Nielen M, van Nes A. 2010. Pleuritis in slaughter pigs: Relations between lung lesions and bacteriology in 10 herds with high pleuritis. Res Vet Sci 88:11–15.

27. Vötsch D, Willenborg M, Weldearegay YB, Valentin-Weigand P. 2018. Streptococcus suis - The “two faces” of a pathobiont in the porcine respiratory tract. Front Microbiol 9:480.

28. Garcia-Graells C, Antoine J, Larsen J, Catry B, Skov R, Denis O. 2012. Livestock veterinarians at high risk of acquiring methicillin-resistant Staphylococcus aureus ST398. Epidemiol Infect 140:383–389.

29. Schulz J, Friese A, Klees S, Tenhagen BA, Fetsch A, Rösler U, Hartung J. 2012. Longitudinal study of the contamination of air and of soil surfaces in the vicinity of pig barns by livestock-associated methicillin-resistant Staphylococcus aureus. Appl Environ Microbiol 78:5666–5671.

30. Bos MEH, Verstappen KM, Van Cleef BAGL, Dohmen W, Dorado-García A, Graveland H, Duim B, Wagenaar JA, Kluytmans JAJW, Heederik DJJ. 2016. Transmission through air as a possible route of exposure for MRSA. J Expo Sci Environ Epidemiol 26:263–269.

31. Pena Cortes LC, LeVeque RM, Funk J, Marsh TL, Mulks MH. 2018. Development of the tonsillar microbiome in pigs from newborn through weaning. BMC Microbiol 18:35.

32. Hwang SM, Kim MS, Park KU, Song J, Kim EC. 2011. tuf gene sequence analysis has greater discriminatory power than 16S rRNA sequence analysis in identification of clinical isolates of coagulase-negative staphylococci. J Clin Microbiol 49:4142–4149.

33. McMurray CL, Hardy KJ, Calus ST, Loman NJ, Hawkey PM. 2016. Staphylococcal species heterogeneity in the nasal microbiome following antibiotic prophylaxis revealed by tuf gene deep sequencing. Microbiome 4:63.

34. Kosecka-Strojek M, Wolska M, Żabicka D, Sadowy E, Międzobrodzki J. 2020. Identification of Clinically Relevant Streptococcus and Enterococcus Species Based on Biochemical Methods and 16S rRNA, sodA, tuf, rpoB, and recA Gene Sequencing. Pathog 2020, Vol 9, Page 939 9:939.

35. Heikens E, Fleer A, Paauw A, Florijn A, Fluit AC. 2005. Comparison of genotypic and phenotypic methods for species-level identification of clinical isolates of coagulase-negative staphylococci. J Clin Microbiol 43:2286–2290.

36. Yatsunenko T, Rey FE, Manary MJ, Trehan I, Dominguez-Bello MG, Contreras M, Magris M, Hidalgo G, Baldassano RN, Anokhin AP, Heath AC, Warner B, Reeder J, Kuczynski J, Caporaso JG, Lozupone CA, Lauber C, Clemente JC, Knights D, Knight R, Gordon JI. 2012. Human gut microbiome viewed across age and geography. Nat 2012 4867402 486:222–227.

37. Schloss PD, Schubert AM, Zackular JP, Iverson KD, Young VB, Petrosino JF. 2012. Stabilization of the murine gut microbiome following weaning. Gut Microbes 3:383.

38. Choudhury R, Middelkoop A, Bolhuis JE, Kleerebezem M. 2019. Legitimate and reliable determination of the age-related intestinal microbiome in young piglets; rectal swabs and fecal samples provide comparable insights. Front Microbiol 10.

39. Choudhury R, Middelkoop A, Boekhorst J, Gerrits WJJ, Kemp B, Bolhuis JE, Kleerebezem M. 2021. Early life feeding accelerates gut microbiome maturation and suppresses acute post-weaning stress in piglets. Environ Microbiol 23:7201–7213.

40. Kumpitsch C, Koskinen K, Schöpf V, Moissl-Eichinger C. 2019. The microbiome of the upper respiratory tract in health and disease. BMC Biol 17:87.

41. Slifierz MJ, Friendship RM, Weese JS. 2015. Longitudinal study of the early-life fecal and nasal microbiotas of the domestic pig. BMC Microbiol 15:184.

42. Fredriksen S, Guan X, Boekhorst J, Molist F, van Baarlen P, Wells JM. 2022. Environmental and maternal factors shaping tonsillar microbiota development in piglets. BMC Microbiol 22.

43. Wang M, Radlowski EC, Monaco MH, Fahey GC, Gaskins HR, Donovan SM. 2013. Mode of delivery and early nutrition modulate microbial colonization and fermentation products in neonatal piglets. J Nutr 143:795–803.

44. Obregon-Gutierrez P, Bonillo-Lopez L, Correa-Fiz F, Sibila M, Segalés J, Kochanowski K, Aragon V, Kochanowski# K, Aragon# V. 2023. Gut-associated microbes are present and active in the pig nasal cavity. bioRxiv 10.1101/2023.06.12.544581.

45. Obregon-Gutierrez P, Aragon V, Correa-Fiz F. 2021. Sow contact is a major driver in the development of the nasal microbiota of piglets. Pathogens 10:697.

46. Correa-Fiz F, Gonçalves dos Santos JM, Illas F, Aragon V. 2019. Antimicrobial removal on piglets promotes health and higher bacterial diversity in the nasal microbiota. Sci Rep 9:6545.

47. Davis M, Dalton K, Johnson Z, Ludwig S, Sabella K, Newman M, Whaley SB, Keet C, McCormack MC, Carroll KC, Matsui EC. 2018. Household Pets and Recovery of Moraxella catarrhalis and Other Respiratory Pathogens From Children With Asthma. Open Forum Infect Dis 5:692–693.

48. McCauley KE, DeMuri G, Lynch K, Fadrosh DW, Santee C, Nagalingam NN, Wald ER, Lynch S V. 2021. Moraxella-dominated pediatric nasopharyngeal microbiota associate with upper respiratory infection and sinusitis. PLoS One 16:e0261179.

49. López-Serrano S, Galofré-Milà N, Costa-Hurtado M, Pérez-De-Rozas AM, Aragon V, Aragon V. 2020. Heterogeneity of Moraxella isolates found in the nasal cavities of piglets. BMC Vet Res 16:1–12.

50. Wang Q, Cai R, Huang A, Wang X, Qu W, Shi L, Li C, Yan H. 2018. Comparison of Oropharyngeal Microbiota in Healthy Piglets and Piglets With Respiratory Disease. Front Microbiol 9:3218.

51. Strube ML, Hansen JE, Rasmussen S, Pedersen K. 2018. A detailed investigation of the porcine skin and nose microbiome using universal and Staphylococcus specific primers. Sci Rep 8:1–9.

52. Taylor DJ. 2006. Pig diseases, 6th ed.

53. Meng F, Wu NH, Nerlich A, Herrler G, Valentin-Weigand P, Seitz M. 2015. Dynamic virus-bacterium interactions in a porcine precision-cut lung slice coinfection model: Swine influenza virus paves the way for Streptococcus suis infection in a two-step process. Infect Immun 83:2806–2815.

54. Fablet C, Marois C, Kuntz-Simon G, Rose N, Dorenlor V, Eono F, Eveno E, Jolly JP, Le Devendec L, Tocqueville V, Quéguiner S, Gorin S, Kobisch M, Madec F. 2011. Longitudinal study of respiratory infection patterns of breeding sows in five farrow-to-finish herds. Vet Microbiol 147:329–339.

55. Jochum L, Stecher B. 2020. Label or Concept – What Is a Pathobiont? Trends Microbiol 28:789–792.

56. Ibarbalz FM, Pérez MV, Figuerola ELM, Erijman L. 2014. The Bias Associated with Amplicon Sequencing Does Not Affect the Quantitative Assessment of Bacterial Community Dynamics. PLoS One 9:e99722.

57. Allali I, Arnold JW, Roach J, Cadenas MB, Butz N, Hassan HM, Koci M, Ballou A, Mendoza M, Ali R, Azcarate-Peril MA. 2017. A comparison of sequencing platforms and bioinformatics pipelines for compositional analysis of the gut microbiome. BMC Microbiol 17:1–16.

58. Wyllie AL, Chu MLJN, Schellens MHB, Gastelaars JVE, Jansen MD, Van Der Ende A, Bogaert D, Sanders EAM, Trzciński K. 2014. Streptococcus pneumoniae in saliva of Dutch primary school children. PLoS One 9:e102045.

59. Barnett DJ m., Arts IC w., Penders J. 2021. microViz: an R package for microbiome data visualization and statistics. J Open Source Softw 6:3201.

60. Friedman J, Alm EJ. 2012. Inferring Correlation Networks from Genomic Survey Data. PLOS Comput Biol 8:e1002687.

61. Van Dongen S, Abreu-Goodger C. 2012. Using MCL to extract clusters from networks. Methods Mol Biol 804:281–295.

